# CALISTA: Clustering and Lineage Inference in Single-Cell Transcriptional Analysis

**DOI:** 10.1101/257550

**Authors:** Nan Papili Gao, Thomas Hartmann, Tao Fang, Rudiyanto Gunawan

## Abstract

We present CALISTA (Clustering and Lineage Inference in Single-Cell Transcriptional Analysis), a numerically efficient and highly scalable toolbox for an end-to-end analysis of single-cell transcriptomic profiles. CALISTA includes four essential single-cell analyses for cell differentiation studies, including single-cell clustering, reconstruction of cell lineage specification, transition gene identification, and pseudotemporal cell ordering. In these analyses, we employ a likelihood-based approach where single-cell mRNA counts are described by a probabilistic distribution function associated with stochastic gene transcriptional bursts and random technical dropout events. We evaluated the performance of CALISTA by analyzing single-cell gene expression datasets from *in silico* simulations and various single-cell transcriptional profiling technologies, comprising a few hundreds to tens of thousands of cells. A comparison with existing single-cell expression analyses, including MONOCLE 2 and SCANPY, demonstrated the superiority of CALISTA in reconstructing cell lineage progression and ordering cells along cell differentiation paths. CALISTA is freely available on https://www.cabselab.com/calista.

## Introduction

The differentiation of stem cells into multiple cell types relies on the dynamic regulation of gene expression (Ralston and Shaw, 2008). In this regard, advances in single-cell gene transcriptional profiling technology have given a tremendous boost in elucidating the decision making process governing stem cell commitment to different cell fates (Kalisky et al., 2018). The applications of single-cell transcriptional analysis have led to new insights on the functional role of cell-to-cell gene expression heterogeneity in the physiological cell differentiation process (Cacchiarelli et al., 2015; Guo et al., 2010; Kumar et al., 2014; Petropoulos et al., 2016; Richard et al., 2016). Along with the surge in single-cell transcriptional profiling studies, algorithms for analyzing single-cell transcriptomics data have received increasing attention. In comparison to measurements from aggregate or bulk samples of cell population, single-cell gene expression profiles display much higher variability, not only due to technical reasons, but also because of the intrinsic stochastic (bursty) dynamics of the gene transcriptional process (Kærn et al., 2005). In particular, the stochastic gene transcription has been shown to generate highly non-Gaussian mRNA count distributions (Raj et al., 2006), which complicate data analysis using established methods that rely on a standard noise distribution model (e.g. Gaussian or Student’s *t*-distribution).

Numerous algorithms have recently been developed specifically for the analysis of single-cell gene expression data. A class of these computational algorithms is geared toward identifying cell groups or clusters within a heterogeneous cell population. Traditional clustering algorithms such as *k*-means and hierarchical clustering have been applied for such a purpose (Grün et al., 2015; Stumpf et al., 2017; Treutlein et al., 2016). Several other single-cell clustering strategies, such as CIDR (Lin et al., 2017), pcaREDUCE (žurauskienė and Yau, 2016) and SNN-cliq (Xu and Su, 2015), adapt more advanced algorithms such as nearest neighbors search. Time-variant clustering strategies have also been implemented to elucidate the appearance of multiple cell lineages (Huang et al., 2014; Marco et al., 2014). In addition, consensus clustering methods, such as SC3 (Kiselev et al., 2017), have received much interest thanks to their superior stability and robustness. Finally, a likelihood-based method called SABEC (Simulated Annealing for Bursty Expression Clustering) (Ezer et al., 2016) employs a mechanistic model of the bursty stochastic dynamics of gene transcriptional process to cluster cells.

Another important class of algorithms deals with the reconstruction of lineage progression during cell differentiation process and the pseudotemporal ordering of single cells along the cell developmental path(s). The lineage progression describes the transition of stem cells through one or several developmental stages during the cell differentiation. This progression may comprise a single developmental path from the progenitor cells to one final cell fate, as well as bifurcating paths leading to multiple cell fates. In this class of algorithms, the reconstruction of the lineage progression and developmental paths is commonly implemented for the purpose of pseudotemporal cell ordering. The pseudotime of a cell represents the relative position of the cell along the developmental path and is typically normalized to be between 0 and 1. By plotting the gene expression against the pseudotime of the cells along a developmental path, one obtains a dynamic trajectory of the gene expression based on which the gene regulations driving the cell fate decision-making can be inferred. Numerous algorithms are available for single-cell transcriptional data analysis for cell lineage inference and cell ordering, notably DPT (Haghverdi et al., 2016), MONOCLE 2 (Qiu et al., 2017; Trapnell et al., 2014) and PAGA (Wolf et al., 2018b) (for a more complete list, see a recent review by Cannoodt et al., 2016).

In this work, we developed CALISTA (Clustering and Lineage Inference in Single Cell Transcriptional Analysis), a numerically efficient and highly scalable toolbox for an end-to-end analysis of single-cell transcriptomics data. CALISTA is capable of and has been tested for analyzing datasets from major single-cell transcriptional profiling technologies, including scRT-qPCR and scRNA-sequencing with both plate-based (e.g. SMART-seq) and droplet-based platforms (scDrop-seq). CALISTA enables four essential analyses of single-cell transcriptomics in stem cell differentiation studies, namely single-cell clustering, reconstruction of lineage progression, transition gene identification and cell pseudotime ordering. In existing literature, these analyses are typically carried out by stringing several task-specific tools together in a bioinformatics pipeline. But, the basic assumptions behind different tools (e.g., regarding the distribution of data noise) maybe incompatible, an issue that has not been delved into more carefully in the literature. In contrast, the different analyses in CALISTA are fully compatible with each other as they are based on the same likelihood-based approach using probabilistic models of gene transcriptional bursts and random dropout events.

In the next section, we describe the algorithmic aspects and functionalities of CALISTA. The single-cell clustering of CALISTA is adapted from a previous method SABEC (Simulated Annealing for Bursty Expression Clustering) with a significant improvement in computational times, while the remaining CALISTA analyses represent novel contributions. For this reason, we focus the performance evaluation of CALISTA on lineage inference and cell pseudotime ordering, and compare CALISTA with widely-used bioinformatics packages including MONOCLE2 (Qiu et al., 2017; Trapnell et al., 2014) and SCANPY (Wolf et al., 2018a). Subsequently, we illustrate CALISTA’s end-to-end analysis using single-cell transcriptional profiles from the differentiation of human induced pluripotent stem cells (iPSCs) into mesodermal (M) or undesired endodermal (En) cells (Bargaje et al., 2017). Finally, we demonstrate the scalability of CALISTA in analyzing large datasets from scDrop-seq studies.

## Results

### Single-cell Transcriptional Analysis using CALISTA

Fig. 1 summarizes the four analyses of single-cell transcriptional profiles in CALISTA, including: (1) clustering of cells, (2) reconstruction of cell lineage progression (3) identification of key transition genes, and (4) pseudotemporal ordering of cells. In CALISTA, we adopt a likelihood-based approach where the likelihood of a cell is computed using a probability distribution of mRNA defined according to the two-state model of gene transcriptional process (Peccoud and Ycart, 1995) and when appropriate, a random dropout event model (see Methods). A random dropout occurs when mRNA molecules of a gene are not detected even though the true mRNA count is non-zero. The single-cell clustering in CALISTA is an adaptation of the algorithm SABEC (Ezer et al., 2016), where the single-cell clustering is carried out in two steps as illustrated in Figure 1b: (1) independent runs of maximum likelihood clustering, and (2) consensus clustering. SABEC has a high computational requirement which hinders its application to large single-cell transcriptomics datasets with 10s-100s of thousands of cells from newer high-throughput single-cell technologies, such as scDrop-Seq. CALISTA offers a substantial numerical speed-up over SABEC thanks to the implementation of a greedy algorithm and the reduction in the model parameter space (see Supplementary Note 1 and Supplementary File S1). CALISTA offers a parallel computing option which enables running the analysis over multiple computing cores for further speed-up.

**Figure 1.**
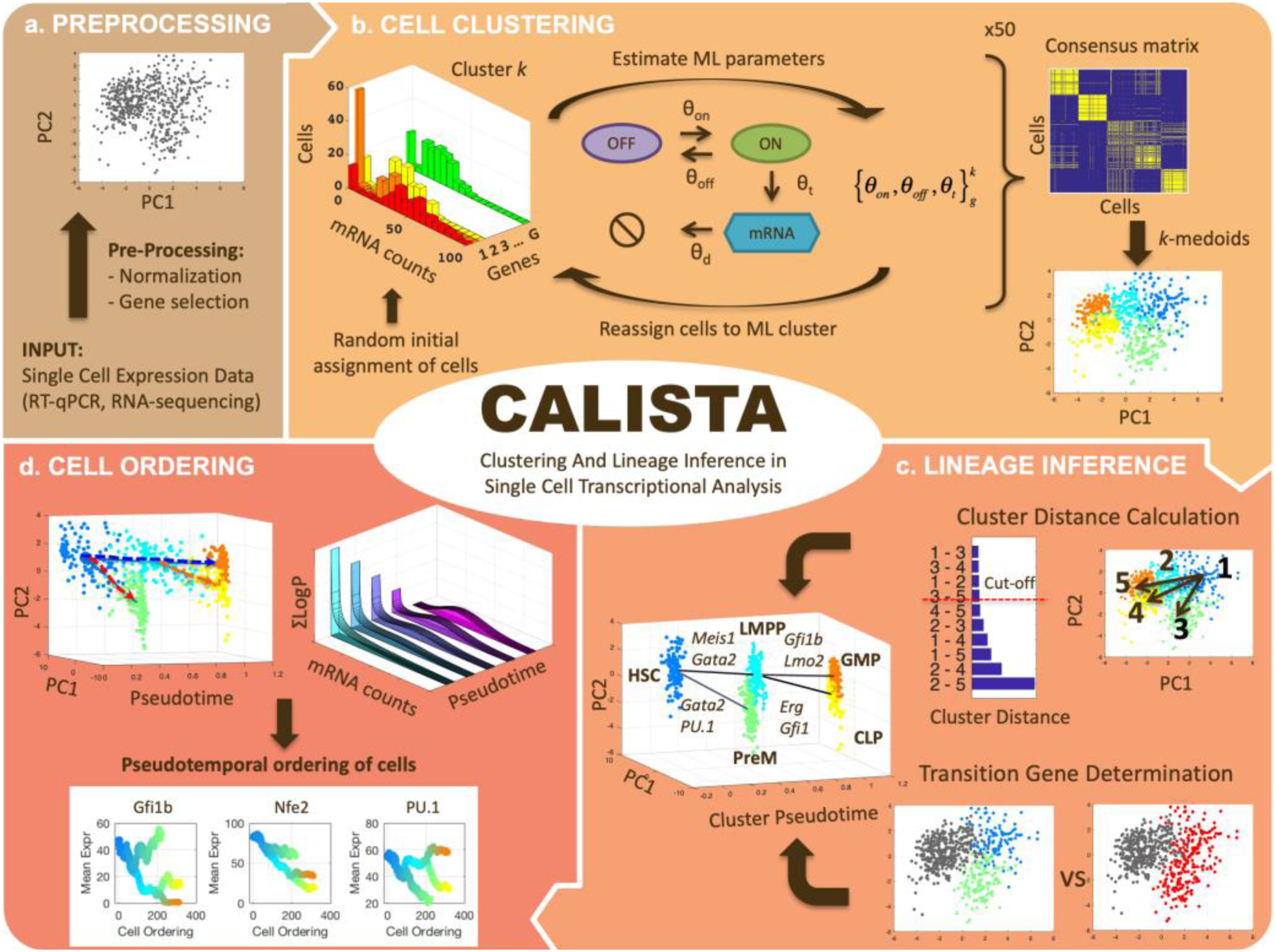
CALISTA single-cell analysis workflow. (a) Single-cell expression input data are first pre-processed. (b) Single-cell clustering in CALISTA combines maximum likelihood and consensus clustering. In the maximum likelihood step, CALISTA relies on the two-state model of the gene transcription process in combination with a model of random dropouts to describe the distribution of single-cell gene expression. The maximum likelihood clustering is implemented multiple times, each starting from random cell assignment into clusters, using a greedy algorithm, the results from which are then used to generate a consensus matrix. The final step implements *k*-medoids clustering algorithm using the consensus matrix. (c) CALISTA uses cluster distances - a measure of dissimilarity in the gene expression distribution between any two clusters – to reconstruct the lineage progression graph. The state transition edges are added in increasing magnitude of cluster distances. In addition, CALISTA provides transition genes between any two connected clusters in the lineage graph based on the gene-wise likelihood differences between having the cells separately and having them in one cluster. (d) For pseudotemporal ordering of the cells, CALISTA reassigns each cell to a transition edge and computes its pseudotime by maximizing the cell likelihood. CALISTA uses a linear interpolation for evaluating the cell likelihood between two connected clusters. Given a developmental path in the lineage progression, CALISTA generates a moving averaged gene expression trajectory using the pseudotemporally ordered cells in the path.

The rest of the single-cell analyses in CALISTA represent new contributions of this work. For reconstructing cell lineage progression, we treat single-cell clusters as cell states through which stem cells transition during the cell differentiation. Here, CALISTA allows the calculation of distances between any pair of cell clusters. The cluster distance – defined as the maximum difference in the cumulative likelihood value upon reassigning the cells from the original cluster to the other cluster (see Methods) – gives a measure of dissimilarity in their gene expression distributions between any two clusters. CALISTA generates the lineage progression graph by sequentially adding state transition edges connecting closely distanced clusters until every cell cluster is connected to at least another cluster (Fig. 1c). CALISTA also provides an interface for users to edit the lineage progression graph, i.e. adding or removing state transition edges, based on the cluster distances and other available information about the cell differentiation. For assigning directionality to the edges, CALISTA relies on user-provided information, for example information on the cell stage or sampling time, the starter/progenitor cells or the expected temporal profiles of the gene expression.

For any two connected clusters in the lineage progression, one can further use CALISTA to obtain the set of transition genes. The transition genes are determined based on the differences of the likelihood between having the cells in separate clusters and having them together in a single cluster (see Methods). Here, the likelihood difference corresponding to a gene reflects the informative power of that gene for segregating cells into two clusters. The transition genes may point to candidate gene markers and genes regulating the state transition during differentiation. SABEC also allows the determination of transition genes, but using a different strategy, called EPiK (Estimation of Pairwise changes in Kinetics), that is based on the statistical significance of the difference in the two-state model parameters between any two clusters.

The final component of CALISTA concerns with the pseudotemporal ordering of cells along a developmental path – defined as a sequence of connected clusters – in the lineage progression graph (Fig. 1d). More specifically, given a developmental path in the reconstructed lineage progression, CALISTA produces a list of the cells ordered in increasing pseudotimes. For this purpose, we first assign a pseudotime to each cluster, which is normalized such that the starting cluster in the lineage progression graph has a pseudotime of 0 and the final cell cluster (or clusters) has a pseudotime of 1. Subsequently, we assign each cell to a transition edge that is pointing to or emanating from the cluster to which the cell belongs, again by adopting the maximum likelihood principle (see Methods). Here, we assume that the distribution of the single-cell gene expression varies monotonically between cell states (clusters). For simplicity, the likelihood of a cell along a transition edge is computed using a linear interpolation of the cell likelihood values from the connected clusters. Each cell is then assigned to the transition edge that maximizes its likelihood value. Analogously, the cell pseudotime is computed by a linear interpolation of the cluster pseudotimes and set to the corresponding maximum point of the cell likelihood value.

### Comparison of CALISTA performance with other methods

We compared the performance of CALISTA with two widely-used single-cell bioinformatics packages for lineage inference and pseudotime cell ordering: MONOCLE 2 (Qiu et al., 2017) and SCANPY (Wolf et al., 2018a). More specifically, in SCANPY package, we used Partition-based Graph Abstraction (PAGA) for lineage progression inference (Wolf et al., 2017) and Diffusion Pseudotime (DPT) for pseudotime cell ordering (Haghverdi et al., 2016).

In the first comparison, we generated *in silico* single-cell expression data of the cell differentiation of central nervous system (CNS) using a stochastic differential equation (SDE) model proposed by Qiu *et al*. (Qiu et al., 2012, n.d.). We simulated single-cell data for 9 time points and 200 cells per time point, totaling 1800 cells (see Methods). As shown in Figure 2a, the simulated single-cell data clearly display two cell lineage bifurcations, as expected in this cell differentiation system (Qiu et al., 2012): (1) CNS precursors (pCNSs) differentiating into neurons and glia cells; (2) glia cells differentiating into astrocytes and oligodendrocytes (ODCs). Figures 2b-d show the reconstructed lineage progressions produced by MONOCLE 2, PAGA, and CALISTA, respectively. PAGA produced the most inaccurate lineage, deviating significantly from the expected lineage (Fig. 2c vs. Fig. 2a). MONOCLE 2 performed better than PAGA, producing a lineage progression that is in general agreement with the *in silico* lineage graph. But, looking at MONOCLE 2’s lineage more carefully, the method identified many more bifurcation or branching points than expected (13 vs. 2). CALISTA outperformed both MONOCLE 2 and PAGA, generating a lineage progression that agrees very well with the *in silico* lineage.

**Figure 2.**
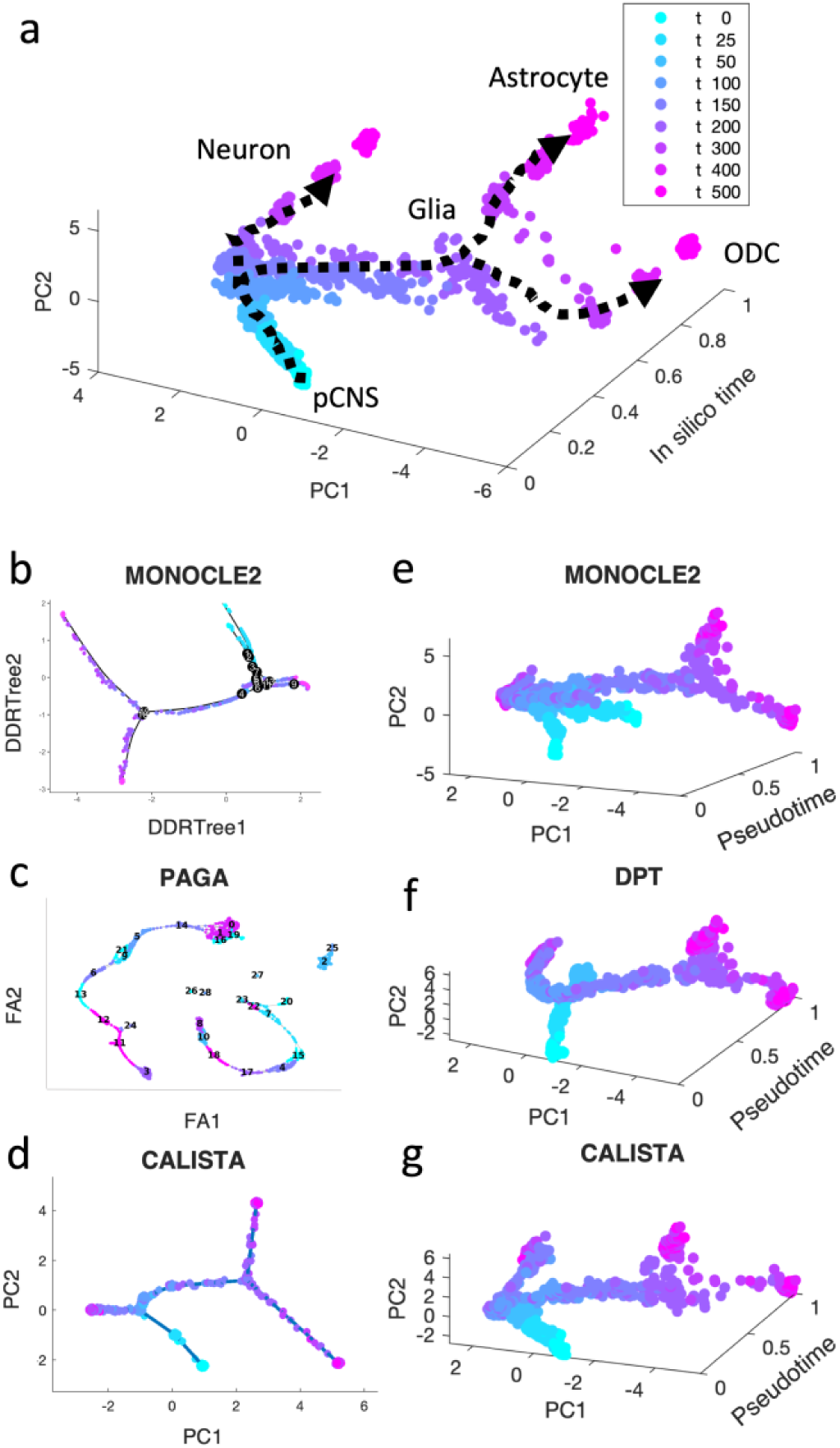
Performance comparison of CALISTA, MONOCLE 2 and SCANPY (PAGA and DPT) using *in silico* single-cell gene expression data of cell differentiation in the central nervous system (CNS). (a) Single-cell gene expression data of CNS differentiation simulated using a model proposed by Qiu *et al*. show two branching / bifurcation points (Qiu et al., 2012): (1) Progenitor CNSs forming neurons and glia cells; (2) Glia cells forming astrocytes and oligodendrocytes (ODCs). (b-d) Reconstructed lineage progression by MONOCLE 2, PAGA (via SCANPY) and CALISTA, respectively. DDRTree: discriminative dimensionality reduction via learning tree (Mao et al., 2015), FA: ForceAtlas2 (Hua et al., 2018), PC: principal component. (e-g) Pseudotemporal ordering of cells by MONOCLE 2, DPT, and CALISTA, respectively.

Figures 2e-f depict the pseudotemporal cell ordering for the simulated CNS single-cell expression produced by MONOCLE2, DPT, and CALISTA, respectively. Besides visual comparisons of the pseudotemporal ordering, we also computed the correlations between the pseudotimes from each of the methods and the *in silico* times of the cells, i.e. the simulation times at which the single-cell mRNA data were sampled (see Supplementary File S2). Among the three algorithms compared, CALISTA’s pseudotimes have the highest correlation with the *in silico* cell times (correlation *ρ* of 0.856), followed by DPT (*ρ = 0.769*) and lastly MONOCLE 2 (*ρ* = 0.571). The cell orderings in Figures 2e-g further confirm the advantage of CALISTA over the other methods.

We further evaluated CALISTA’s performance using four single-cell gene transcriptional datasets from cell differentiation systems with a variety of lineage topologies, including Bargaje et al. study on the differentiation of human induced pluripotent stem cells (iPSC) into cardiomyocytes (Bargaje et al., 2017), Chu et al. study on the differentiation of human embryonic stem cells (hESC) into endodermal cells (Chu et al., 2016), Moignard et al. study on hematopoietic stem cell (HSC) differentiation (Moignard et al., 2013), and Treutlein et al. study on mouse embryonic fibroblast (mEF) differentiation into neurons (Treutlein et al., 2016). We again compared CALISTA with the same single-cell analyses as in the *in silico* case study above. Figures 3 summarizes the reconstructed lineage progression of the cell differentiation using MONOCLE 2, PAGA, and CALISTA. The cell differentiation in these cell systems follows the lineage progression drawn in Figure 4a. As in the *in silico* case study above, CALISTA generated the most accurate lineage progressions, followed by MONOCLE 2 and lastly PAGA. Figures 4b-c show the pseudotemporal ordering of cells produced by MONOCLE 2, DPT, and CALISTA, respectively. In assessing the accuracy of the pseudotimes, we relied on the known lineage progression and cell capture times, since the true cell differentiation times are not known in these datasets. All three methods performed equally well for the HSC differentiation dataset by Moignard et al. (Moignard et al., 2013). For iPSC dataset (Bargaje et al., 2017), CALISTA gave the most accurate pseudotimes, while MONOCLE2 and DPT had difficulties in assigning pseudotimes for one of the final cell type due to the close similarity of the gene expression with the progenitor iPSCs (see next section for more detail). For mEF dataset (Treutlein et al., 2016), CALISTA produced pseudotimes that are most consistent with the known lineage and capture times, followed by DPT and then MONOCLE 2. Finally, for hESC dataset (Chu et al., 2016), CALISTA again outperformed DPT and MONOCLE2, but here MONOCLE 2 performed better than DPT. In summary, for both simulated and real life single-cell gene expression datasets, CALISTA is able to reconstruct lineage progression and single-cell pseudotimes much better than widely-used single-cell gene expression analyses.

**Figure 3.**
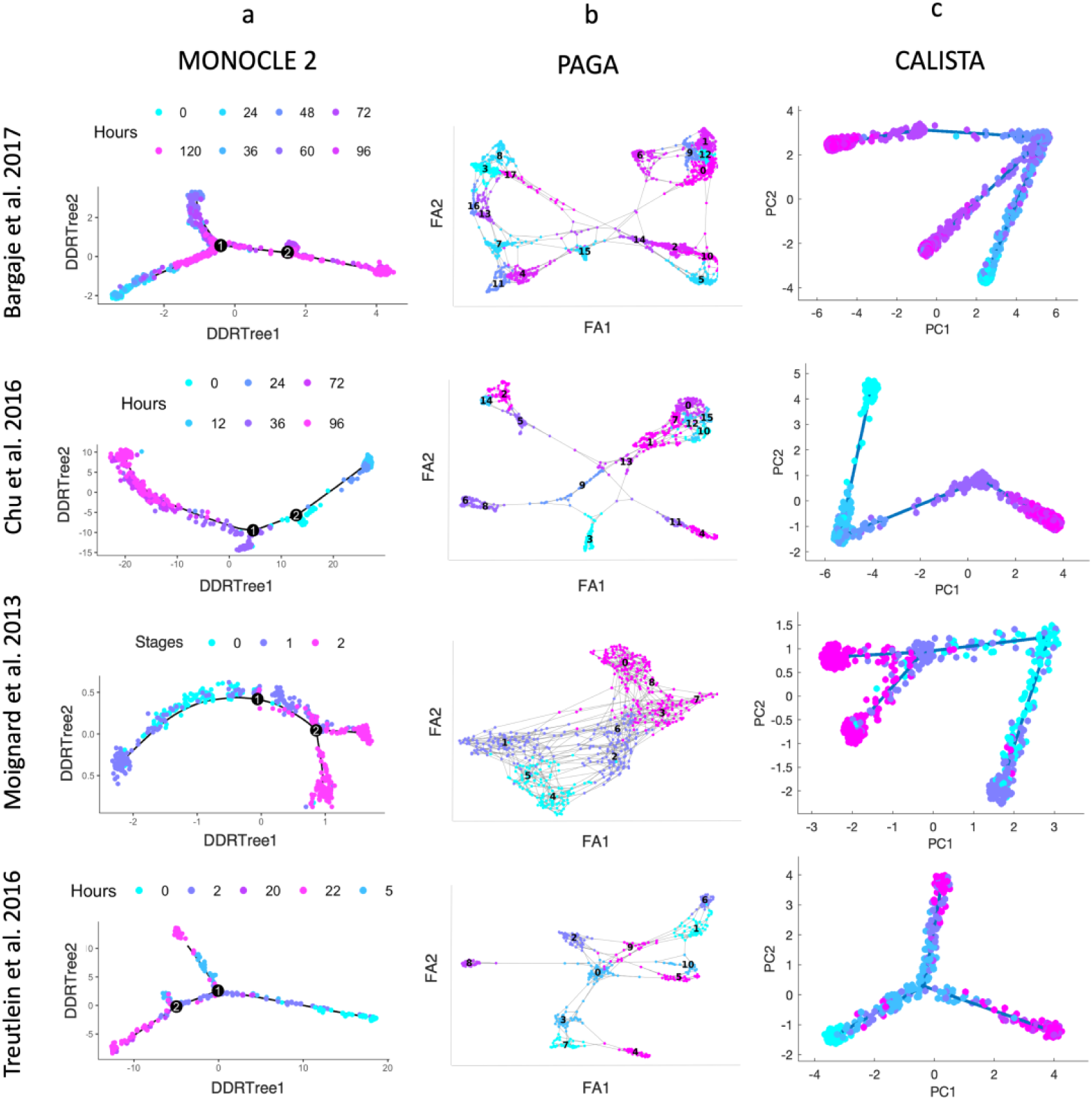
Comparison of lineage progressions reconstructed from single-cell transcriptional profiles by MONOCLE 2, PAGA, and CALISTA. (Top row) Induced pluripotent stem cell (iPSC) differentiation into cardiomyocytes in Bargaje et al. study (Bargaje et al., 2017). (Second row) Human embryonic stem cell differentiation into endodermal cells in Chu et al. study (Chu et al., 2016). (Third row) Hematopoietic stem cell differentiation in Moignard et al. study (Moignard et al., 2013). (Bottom row) Mouse embryonic fibroblast differentiation into neurons in Treutlein et al. study (Treutlein et al., 2016). (Left column) MONOCLE 2. (Middle column) PAGA. (Right column) CALISTA. DDRTree: discriminative dimensionality reduction via learning tree (Mao et al., 2015), FA: ForceAtlas2 (Hua et al., 2018), PC: principal component. The colors indicate the cell sampling times.

**Figure 4.**
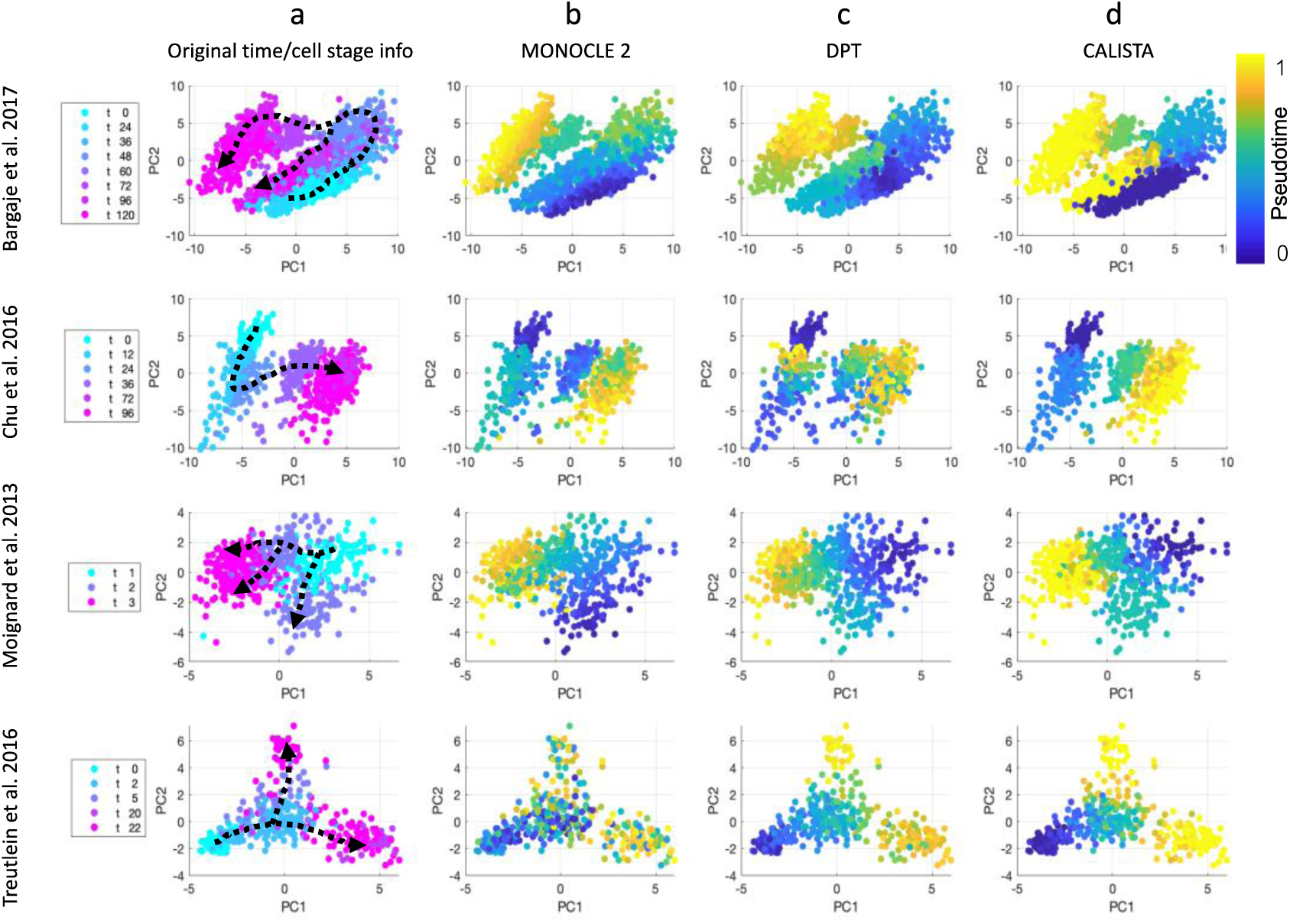
Comparison of pseudotemporal cell ordering using single-cell transcriptional profiles by MONOCLE 2, DPT, and CALISTA. (Top row) Induced pluripotent stem cell (iPSC) differentiation into cardiomyocytes in Bargaje et al. study (Bargaje et al., 2017). (Second row) Human embryonic stem cell differentiation into endodermal cells in Chu et al. study (Chu et al., 2016). (Third row) Hematopoietic stem cell differentiation in Moignard et al. study (Moignard et al., 2013). (Bottom row) Mouse embryonic fibroblast differentiation into neurons in Treutlein et al. study (Treutlein et al., 2016). (First column) Single-cell gene expression data. Pseudotemporal cell ordering from (Second column) MONOCLE 2, (Third column) DPT, and (Fourth column) CALISTA. PC: principal component. The colors in the first column indicate the cell sampling times, and those in the second-fourth column indicate the pseudotimes.

### Application to the differentiation of induced pluripotent stem cells to cardiomyocytes

In the following, we demonstrated an end-to-end analysis of single-cell gene expression data using CALISTA. Here, we used the single-cell gene expression dataset from the differentiation of human iPSCs into cardiomyocytes in Bargaje et al. study (Bargaje et al., 2017). The dataset includes single-cell expression of 96 genes measured by RT-qPCR for 1896 cells collected across 8 time points (day 0, 1, 1.5, 2, 2.5, 3, 4, 5) after induction to differentiate. Figure 5a shows the developmental states identified in the original study: epiblast cells (E) in the early stage (day 0, 1, 1.5), primitive streak (PS)-like progenitor cells in the intermediate stage (day 2, 2.5), and a lineage bifurcation into either the desired mesodermal (M) or undesired endodermal (En) cell fate in the late stage (day 3, 4, 5).

**Figure 5.**
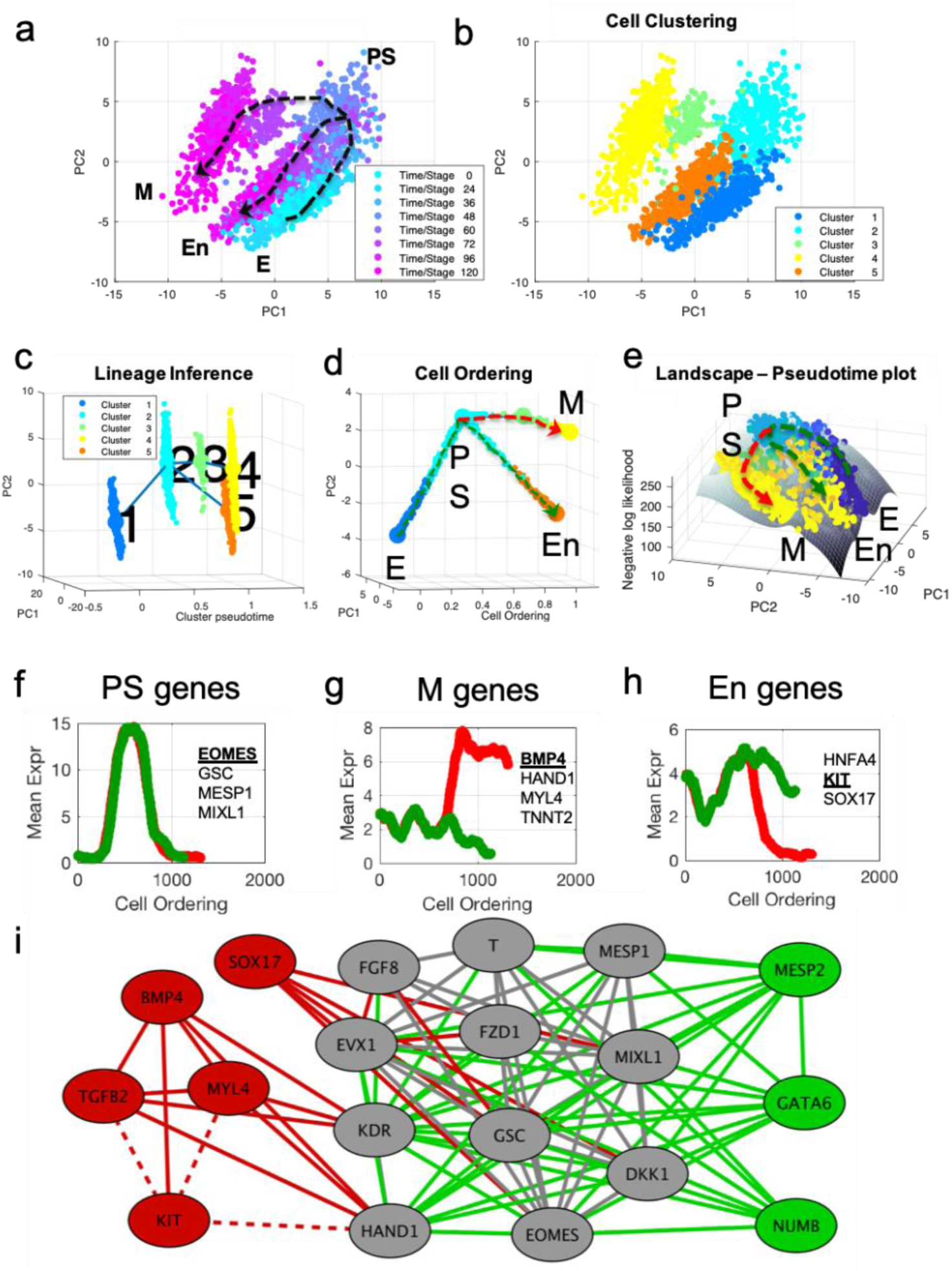
End-to-end analysis of single-cell transcriptional profiles during iPSC differentiation into ardiomyocyte. The single-cell gene expression dataset was taken from the study of Bargaje et al. (Bargaje et al., 2017) (a-b) PCA plots of single-cell dataset along the first two principal components. Each dot represents a cell, where the colors indicate (a) the capture time info and (b) CALISTA single-cell clusters. (c) Lineage progression graph reconstructed by CALISTA. The graph was generated by adding transition edges in increasing order of cluster distances until each cluster is connected by an edge. Following this procedure, an edge connecting cluster 1 and 5 was originally added in the graph. We manually removed this edge as the edge bypassed intermediate capture times. (d) Pseudotemporal ordering of cells. Cells were first assigned to the state transition edges in the lineage progression, and then pseudotemporally ordered along two developmental paths – mesodermal (M) path (red) and endodermal (En) path (green). (e) Cell likelihood landscape. The landscape shows the surface of the negative log-likelihood of the cells. A higher value thus indicates a state of increased gene expression uncertainty. (f-h) Moving window average of gene expression along M and En developmental paths for representative (g) PS, (h) M, and (i) En genes (underlined and boldfaced). (i) Gene modules of M (red and grey) and En paths (green and grey) (drawn using Cytoscape (Smoot et al., 2011)). The grey nodes and edges are common between the M and En modules. Solid (dashed) lines indicate positive (negative) gene correlations.

First, we clustered the cells by using CALISTA. The optimal number of clusters was chosen to be five based on the eigengap plot (see Supplementary Figure S1). The single-cell clustering of CALISTA, as shown in Figure 5b, recapitulates the previously identified developmental states. Here, clusters 1, 2, and 5 contain mostly E, PS, and En cells, respectively, while M cells are split between clusters 3 and 4 that are demarcated by different capture times (see cluster compositions in Supplementary Figure S2).

After single-cell clustering, we employed CALISTA to infer the lineage progression graph. The cluster pseudotimes were set to the modes (most frequent values) of the cell capture times in the clusters divided by the maximum cell capture time (see Supplementary Figure S3). The directionality of the state transition edges was set according to the cluster pseudotimes, pointing from a cluster with a lower pseudotime to that with a higher pseudotime. As shown in Figure 5c, the lineage progression graph reconstructed by CALISTA reproduces the lineage bifurcation event as the cells transition from PS-like cells to take on either M or En cell fates (Bargaje et al., 2017). Subsequently, for each state transition edge, we identified the set of transition genes (see Supplementary Figure S4). Among the identified transition genes in the lineage progression graph (25 genes in total, see Supplementary File S3), many are known lineage specific transcriptional regulators involved in the iPSC cell differentiation, such as EOMES, GATA4, GSC, HAND1, KIT, MESP1, SOX17, and T (Bargaje et al., 2017).

Finally, we employed CALISTA to generate the pseudotemporal ordering of cells along the two distinct developmental paths in the lineage progression: (1) the M path forming mesodermal cells (cluster 1 – 2 – 3 – 4; see red dashed path in Figure 5d) and (2) the En path forming endodermal cells (cluster 1 – 2 – 5; see green dashed path in Figure 5d). After assigning cells to the state transition edges and prescribing the cell pseudotimes, we ordered cells belonging to each developmental trajectory in increasing pseudotimes with a total of 1408 cells in the M path and 1215 cells in the En path.

The likelihood value of each cell computed during the pseudotemporal cell ordering can further be visualized as a landscape plot. Figure 5e depicts the negative log-likelihood surface of the cells over the first two principal components. A higher value on the surface indicates a cell state with broader mRNA distributions, i.e. a state of higher uncertainty in the gene expression. As shown in Figure 5e, iPSC cells start their journey from a valley in this surface, implying that the progenitor cells are at a low uncertainty state. As the cell differentiation progresses, cells pass through an intermediate state with higher uncertainty, where a peak uncertainty is reached at or around the cell lineage bifurcation. After the bifurcation, cells follow two paths toward lower uncertainty, leading to two valleys corresponding to distinct cell fates (M and En fates). The rise-and-fall in gene expression uncertainty have also been reported in other cell differentiation systems, suggesting that stem cells go through a transition state of high uncertainty before committing to their final cell fate(s) (Richard et al., 2016; Stumpf et al., 2017).

To visualize the gene expression trajectories along the two cell differentiation paths, we calculated the moving average expression values of transition genes for the pseudotemporally ordered cells using a moving window comprising 10% of the total cells in each path (see Figure 5f-h and Supplementary Figure S5). A number of transition genes follow highly similar expression trajectories along the M and En paths, with an increase in expression from E to PS-like state, followed by a decrease in expression from PS-like to M or En state (see Figure 5f). The majority of genes with the aforementioned trajectory are known PS-like markers (for example EOMES, GSC, MESP1, and MIXL1 (Bargaje et al., 2017; Ng et al., 2005; Tiyaboonchai et al., 2017)), and thus, we refer to these genes as PS-genes (see Supplementary File S3). Another group of transition genes show differential profiles between the M and En paths after the lineage bifurcation. Here, we define M-genes as the genes with higher expressions along the M path than the En path (see Figure 5g). Correspondingly, we define En-genes as genes with higher expressions along the En path than the M path (see Figure 5h). Notably, many of the known M marker genes (e.g., BMP4, HAND1, MYL4 and TNNT2 (Bargaje et al., 2017; Jagtap et al., 2011)) are among the M-genes, and several of the known En marker genes (e.g. HNFA4, KIT, and SOX17 (Bargaje et al., 2017; Ng et al., 2010; Thomas et al., 2006)) are among the En-genes (see Supplementary File S3). In general, after the lineage bifurcation, the expressions of M genes increase along the M path, but are either suppressed (BMP4, HAND1, KDR) or unchanged (MYL4, TGFB2, TNNT2) along the En path. Among the known En markers, the expression of HNFA4 increases along the En path after the lineage bifurcation but remains relatively unchanged along the M path. The expression of KIT follows the opposite profile, where the gene expression is downregulated along the M path and is relatively unchanged along the En path. While the expression of SOX17, like KIT, decreases along the M path, the gene is upregulated along the En path immediately after the lineage bifurcation before being downregulated toward the end of the developmental trajectory.

Finally, we constructed gene co-expression networks for the M and En developmental paths based on the pseudotemporal profiles of the gene expression (pairwise Pearson correlation, *p*-value ≤ 0.01 and correlation value ≥ 0.8, see Supplementary Figure S6). We identified cliques in the M and En gene co-expression networks, i.e. a subset of genes (at least 5) that are connected to each other, using a maximal clique analysis by the Bron-Kerbosch algorithm (Bron and Kerbosch, 1973). Figure 5i depicts the cliques from the M and En gene co-expression networks, showing three gene regulatory modules: one module specific to the M path (red), another specific to the En path (green), and a shared module (grey). Most of the PS genes, including known PS marker genes, belong to the shared module as expected. Among the genes in the shared module, DKK1, FZD1 and T are involved in Wnt signaling pathway, which is known to promote cell differentiation (Davidson et al., 2012). GATA6, which plays an important role in the endoderm commitment (Tiyaboonchai et al., 2017), belongs to the En module. On the other hand, the M module shows activating relationships among mesoderm gene markers (e.g. BMP4, MYL4 and HAND1), and antagonistic relationships between several M genes and KIT, an En gene marker.

### Application of CALISTA to massively parallel Drop-Seq datasets

To demonstrate the scalability of CALISTA, we analyzed single-cell expression datasets from droplet-based assays. Single-cell Drop-seq is a massively parallel genome-wide expression profiling technology capable of analyzing thousands of cells in a single experiment. However, the bioinformatic analysis of large single-cell transcriptomics datasets poses a significant computational challenge (Angerer et al., 2017). To address this challenge, CALISTA offers a parallelization option for handling large datasets by splitting the greedy optimization runs among multiple computing cores (see Supplementary Note S2 for more details on CALISTA implementation).

We first tested the single-cell clustering performance of CALISTA in analyzing Drop-seq datasets using the single-cell study of mouse spinal cord neurons by Sathyamurthy et al. (~18K nuclei) (Sathyamurthy et al., 2018) and the study of peripherical blood mononuclear cells by Zheng et al. (~68K cells) (Zheng et al., 2017). We noted that *k*-medoid clustering for large consensus matrices is computationally prohibitive. Thus, we bypassed the consensus clustering and took among the independent runs of the greedy algorithm, the cell clustering corresponding to the highest likelihood value. For Sathyamurthy et al. dataset, CALISTA clustering identified nine single-cell clusters based on the eigengap plot, which agrees well with the original study (by comparing Supplementary Figure S7 with Figure 1D in the original publication (Sathyamurthy et al., 2018)). For the largest cluster (52% of total data) containing neurons, CALISTA further split the cells into two subpopulations – one comprising cells with high expressions of Snap25, Syp and Rbiox3, and the other comprising cells with medium expression of these genes. Notably, CALISTA was able to distinguish clearly Schwann and Meningeal cells, which in the original study (using SC3 (Kiselev et al., 2017)), were placed into the same cluster (Sathyamurthy et al., 2018). Meanwhile, for Zheng et al. scDrop-seq dataset, CALISTA generated cell clusters that show a general agreement with the original study (see Supplementary Figure S8).

We then tested CALISTA’s lineage progression reconstruction on scDrop-seq data of ~38K cells, taken from 12 developmental stages of zebrafish embryogenesis (Farrell et al., 2018). The results of CALISTA are summarized in Figure 6. For this large dataset, we employed a modified clustering procedure (see Supplementary Note S2) and identified 82 clusters, 23 of which comprise cells at the final developmental stage. Figure 6a depicts the reconstructed lineage progression by CALISTA (see also Supplementary File S5), while Figure 6b shows the cell type labels for the cell clusters at the final developmental stage based on the expression levels of 26 key gene markers (see Supplementary Figure S9). The results are again in good agreement with the known cell types (Farrell et al., 2018).

**Figure 6.**
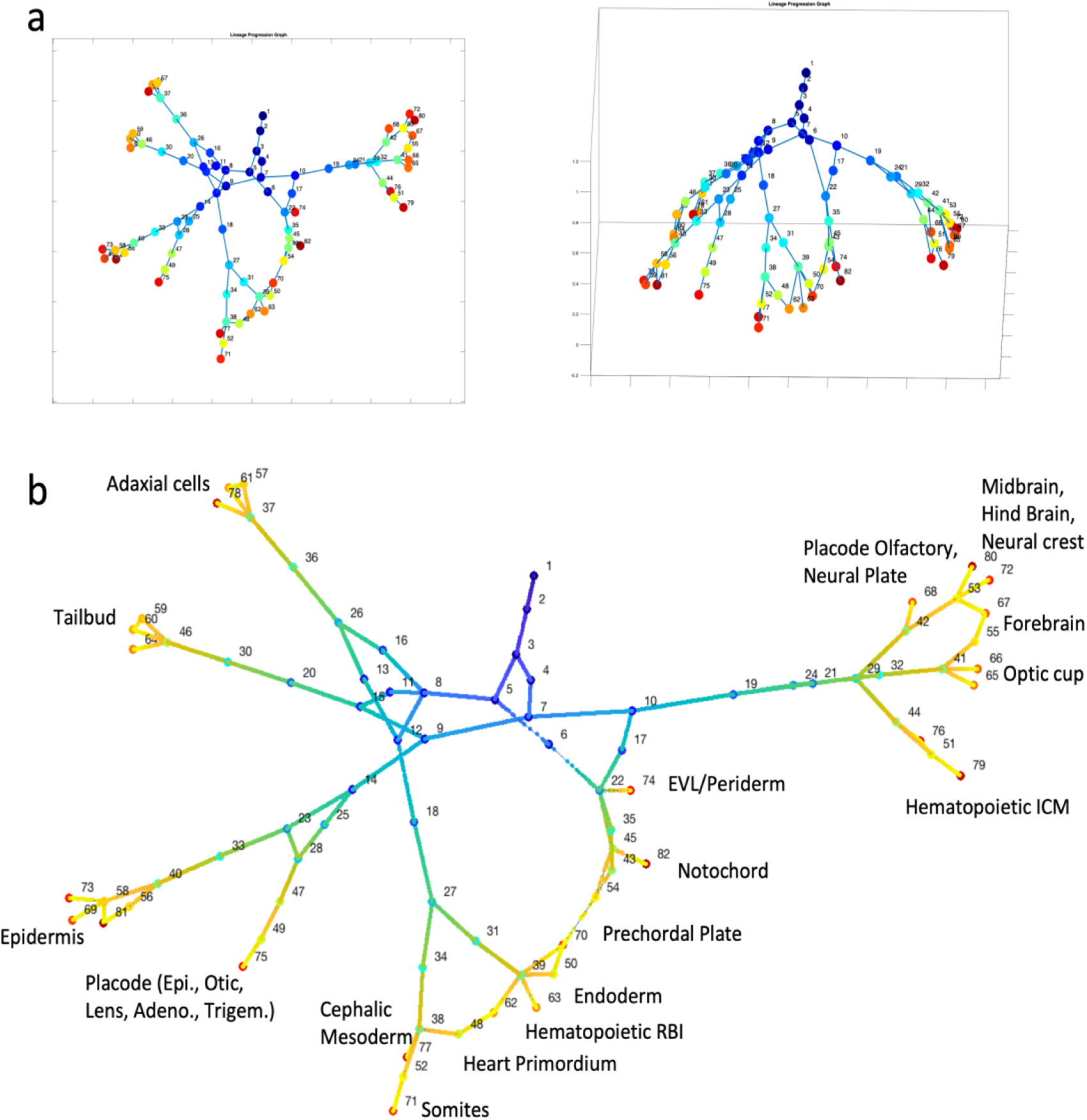
Analysis of single-cell Drop-seq gene expression data of zebrafish embryogenesis. (Farrell et al., 2018). (a) Reconstruction of lineage progression and pseudotimes calculation of cells (see also Supplementary File S5). (b) Cell types at the final developmental stage.

## Discussion

CALISTA provides four bioinformatic analyses for single-cell expression data that are essential in studies of stem cell differentiation. The analyses can be implemented either sequentially in a data analytics pipeline or separately in a standalone application. Throughout the development of the analyses in CALISTA, we used the same likelihood approach based on the two-state stochastic gene transcriptional model. Thus, the analyses are fully compatible with each other. The use of a mechanistic model in CALISTA brings an additional advantage, because the model parameters have relevance to the mechanism of gene transcription process and thus can provide insights into the gene regulations during stem cell differentiation. To the best of our knowledge, CALISTA is the first mechanistic model-based toolbox that allows an end-to-end analysis of single-cell transcriptional profiles. Despite the focus on stem cells in our work, each of the tools in CALISTA is agnostic to the source of the single-cell transcriptomic data analysis and thus can be used for other studies.

CALISTA’s single-cell clustering analysis is an adaptation of another method called SABEC (Ezer et al., 2016), with a significant improvement in computational efficiency, thanks to the implementation of greedy algorithm. This improvement is critical and necessary for the analysis of extremely large single-cell datasets with tens to hundreds of thousands of cells, which will become more common in the near future. In addition to computational efficiency, CALISTA is able to account for random technical dropout events, a feature that is missing from SABEC. With the exception of the single-cell clustering analysis, CALISTA includes novel algorithms for transition gene identification, lineage progression inference and pseudotime calculations. As demonstrated in numerous case studies above, when compared to popular single-cell analysis methods for lineage inference and pseudotemporal cell ordering, including MONOCLE 2, PAGA, and DPT, CALISTA performed better than these methods.

CALISTA further features an interactive interface for user inputs at different steps in the analysis, for example in setting the number of cell clusters or during the curation of the lineage progression graph (see CALISTA tutorials on https://www.cabselab.com/calista). The user interface enables incorporating existing biological knowledge of the cell differentiation system, which is often difficult – if not impossible – to codify. Although such prior knowledge is not necessary for using CALISTA, the ability to incorporate this knowledge, whenever available, is a useful and important feature in the analysis of single-cell transcriptional profiles.

## Declarations

### Availability of data and materials

A MATLAB and R version of CALISTA used in this study is freely available from the following website: https://www.cabselab.com/calista.

All the public single cell data sets analyzed in this study are available from the original publications. *In silico* datasets are available on CALISTA website: https://www.cabselab.com/calista.

### Authors’ contributions

NPG and RG designed the computational framework and workflow and wrote the manuscript; NPG developed open-course tool and performed all data analyses; NPG, TH and TF collected all necessary data and performed the preliminary analysis; All authors read and approved the final manuscript.

### Consent for publication

Not applicable.

## Methods

### Input data and data preprocessing

The single-cell gene expression matrix should be formatted into an *N*×*G* matrix, where *G* denotes the number of genes, *N* denotes the number of cells, and the matrix element *m*_*n*,*g*_ is the transcriptional expression value of gene *g* in the *n*-th cell. CALISTA accepts expression values from RT-qPCR (2^*Ct*^ value) and scRNA-Seq including both plate-based (e.g. log(RPKM) or log(TPM)) and droplet-based measurements (e.g. gene UMI counts). For UMI data from scDrop-seq, CALISTA further scales the expression matrix by dividing each gene UMI count with the total UMI count in the corresponding cell and then multiplying the value with the median of the total UMI counts among cells (Zheng et al., 2017).

Before performing single-cell analysis, we first preprocess the single-cell expression matrix by removing the genes and cells (i.e., columns and rows of the expression matrix, respectively) with a large fraction of zero expression values, exceeding a user-defined threshold (default threshold: 100% for genes and 100% for cells). For scRNA-seq datasets, CALISTA further selects a number of informative genes *Y* for the single-cell analysis following a previously described procedure (Macosko et al., 2015), with *Y* is to the minimum among the following: 200, the number of genes *G*, and a user-defined percentage of the number of cells *N*.

Next, CALISTA scales the single-cell expression values such that the maximum value of any gene is 200. The scaling is carried out as follows:

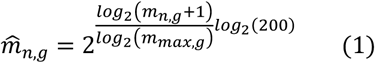

where *m*_*n*,*g*_ and 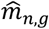 represent the original and scaled expression value of gene *g* in the *n*-th cell respectively, and *m*_*max*,*g*_ is the maximum expression value in gene *g* (i.e. the maximum of *m*_*n*,*g*_ over all cells). The scaling above allows CALISTA to use a pre-computed table for the maximum likelihood in the clustering analysis, thereby reducing the computational cost significantly. We tested using *in silico* single-cell gene expression datasets generated using the two-state gene transcriptional model, and confirmed that the scaling does not affect the clustering accuracy (see Supplementary Note S3). After scaling, for scDrop-Seq data, we re-rank the top *Y* genes in increasing value of the gene-wise likelihood *v*^*g*^ computed following Equation (10) below, with the random dropout model in Equation (4) incorporated in the calculation of likelihood. A lower gene-wise likelihood value indicates a broader distribution of single-cell expression. Genes with likelihood values exceeding a given threshold (by default set at the elbow of the curve of likelihood vs gene rank) is removed from further analysis.

### Stochastic two-state gene transcriptional model

For describing the mRNA distribution of a gene, CALISTA relies on the two-state model developed by Peccoud and Ycart (Peccoud and Ycart, 1995). The model characterizes the stochastic bursty gene transcriptional process at the single-cell level. More specifically, the model describes a promoter that switches stochastically between an OFF (inactive or non-permissive) state, where the gene transcription cannot start, and an ON (active or permissive) state, where the gene transcription can proceed. The set of reactions describing the stochastic gene transcription in the two-state model are as follows:

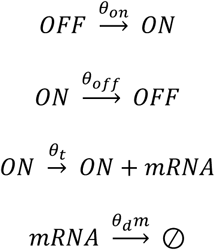

where *θ_on_* is the rate of the promoter activation, *θ_off_* is the rate of the promoter inactivation, *θ_t_* is the rate of mRNA production when the promoter is active, *θ_d_* is the rate constant of mRNA degradation, and *m* denotes the number of mRNA molecules. At steady state, the probability distribution of mRNA count *m* can be approximated by the following density function (Raj et al., 2006):

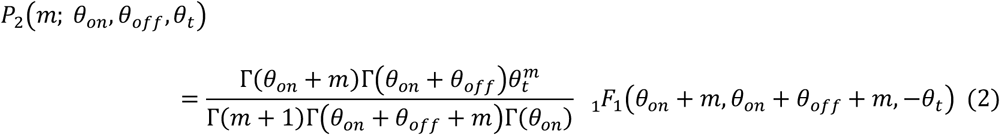

where _1_*F*_1_ represents the confluent hypergeometric function of the first kind.

### Random dropout event model

If desired and when appropriate, users can account for random dropout events in CALISTA. The inclusion of random dropouts is particularly suitable when dealing with single-cell transcriptional profiles from Drop-Seq technology. In CALISTA, the dropout probability is modeled by a negative exponential function with an optimal decay constant of *λ*, as follows:

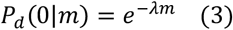

where *P*(0|*m*) denotes the probability that the measured single-cell mRNA count is zero when the true number of mRNA molecules is *m*. A decay constant *λ* = 0 gives a dropout probability of 1, i.e. dropout occurs regardless of the true mRNA count. The parameter *λ* is estimated from the plot of the fraction of zeros against the mean expression across all measured genes, following the procedure described in Pierson et al. (Pierson and Yau, 2015) (see examples in Supplementary Figure S10). Combining the dropout event model with the two-state gene transcription model above give the following distribution function of measured single-cell mRNA readouts 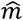

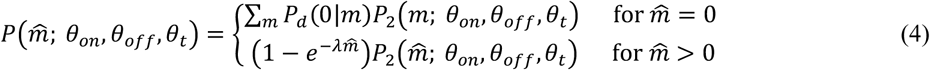

### Cell Clustering

As illustrated in Figure 1b, CALISTA combines maximum likelihood and consensus clustering algorithms for single-cell clustering analysis. Following a previous method SABEC (Ezer et al., 2016), CALISTA uses the stochastic two-state model of gene transcription to derive the steady-state probability distribution of gene expression values. The probability distribution is used to prescribe a likelihood value to a cell based on the measured single-cell expression values of its genes (Ezer et al., 2016). In contrast to SABEC that employs a simulated annealing algorithm, CALISTA implements a greedy algorithm to solve the likelihood maximization problem (see Supplementary Note S1). The greedy algorithm is implemented repeatedly for a specified number of times, each starting from a different random initial cell assignment, to generate a consensus matrix. The (*i*,*j*)-th element of the consensus matrix records the number of times that the *i*-th and *j*-th cells are assigned to a cluster. The final cell clustering is obtained by applying *k*-medoids on the consensus matrix (Bhat, 2014) (see Supplementary Note S1 for more details).

#### Lineage Inference

The first novel algorithm in CALISTA is the reconstruction of cell lineage graph, which reflects the lineage progression in the differentiation process (Figure 1c). Based on the view that cell clusters represent cell states, the nodes of the lineage graph comprise cell clusters, while the edges represent state transitions in the lineage progression. For inferring the lineage graph, CALISTA computes cluster distances based on dissimilarities in the gene expressions among cells from two clusters. Again, CALISTA adopts a likelihood-based strategy using the probability distribution of mRNA from the stochastic two-state gene transcription model to define the cluster distances.

### Cluster distance

Given *K* clusters from CALISTA single-cell clustering analysis above or the clustering provided by the user, CALISTA evaluates a *K* × *K* dissimilarity matrix *S* where the element s*_kj_* gives the likelihood of cells from cluster *k* to be assigned to cluster *j*, computed as follows:

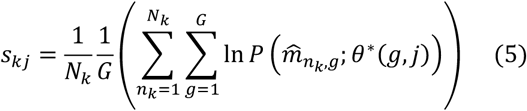

where *N*_*k*_ is the set of cell indices for the cells in cluster *k* and 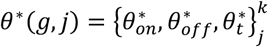 is the parameter vector of two-state gene transcription that maximizes the joint probability of obtaining the gene expression values for gene *g* of the cells in the cluster *j*, as follows:

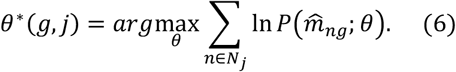

The probability 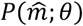 is computed using the steady state probability distribution from the two-state gene transcription model as given in Eq. (2) (assuming that random dropout events are insignificant) or using the distribution function defined in Eq. (4).

Note that the diagonal element s*_kk_* is the sum of the cell likelihood in the original cluster assignment, i.e. 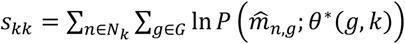, and thus are larger than other elements s*_kj_*, *j* ≠ *k* The dissimilarity coefficients in the matrix S is subsequently normalized by subtracting each element with the diagonal element of the corresponding rows, as follows:

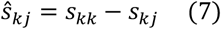

Since the likelihood always takes on negative values, the normalized dissimilarity coefficient assumes a positive value. A larger *Ŝ*_*kj*_ reflects a higher degree of dissimilarity between the two clusters. The distance between the *j*-th and *k*-th cluster, denoted by *d_kj_*, is defined as:

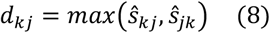

### Lineage graph construction

CALISTA generates a lineage graph by connecting single-cell clusters (states) based on the cluster distances. The lineage graph describes the state transition of stem cells during the cell differentiation process, under the assumption that the transitions occur between closely related cell states (i.e. between clusters with low distances). Briefly, CALISTA starts with a fully disconnected graph of single cell clusters, and sequentially adds one transition edge at a time in increasing magnitude of cluster distance until each cluster is connected by at least one edge. Once the lineage graph has been established, CALISTA assigns directionalities to the edges in the lineage graph according to user-provided information, e.g. starting cells/clusters or expected gene expression profiles. Given information of the starting cells, CALISTA defines the cell cluster(s) containing these cells as the starting cluster(s). On the other hand, given the expected trajectory of some marker genes, CALISTA uses the mean expression of the gene marker(s) in each cell cluster to determine the starting cell cluster.

When the stage or time information is provided for the cells, CALISTA implements the following lineage reconstruction procedure. First, large outliers in the cluster distances (i.e. cluster distances that are larger than the median value by 3 scaled median absolute deviation) are removed from further consideration. Single-cell clusters are then labelled by their most frequent (mode) cell stage or time. Clusters with the lowest stage/time label are the starting clusters. CALISTA constructs a connected graph by assigning one (and only one) incoming edge for each cluster, except for the starting cluster(s), from the cluster with the lowest cluster distance among the set of feasible parent cluster(s). Here, the feasible parents of a given cluster *j* are any clusters with time/stage labels that are the nearest to but do not exceed a cutoff cell times/stages in cluster *j*. The default cutoff is set to the 5^th^ percentile of the cell times/stages in cluster *j*. This cutoff is used to ensure that there exists sufficient difference in the cell times between the parent and daughter cell clusters.

Besides the automated lineage reconstruction above, CALISTA further allows the users to manually add or remove transition edges between pairs of clusters/nodes through a user-friendly GUI (see CALISTA user manual).

#### Transition genes

Another novel contribution in CALISTA is an algorithm to extract the set of transition genes between any two connected clusters in the lineage graph. Here, transition genes are defined as genes whose single-cell expressions are highly informative in grouping cells into the two clusters. More specifically, CALISTA evaluates the likelihood difference between having cells assigned to two separate clusters and having the cells together in one cluster, again using the steady-state distribution of mRNA from the two-state gene transcription model. Given two clusters *j* and *k*, we compute the following:

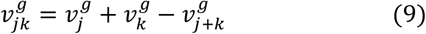

With

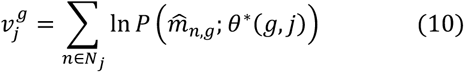

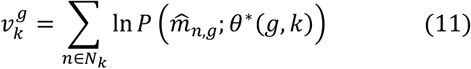

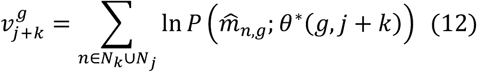

where the optimal parameter vector *θ*^∗^(*g*, *j* + *k*) is obtained according to Equation (6) for all cells from clusters *j* and *k* together. The value of 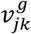 reflects the informativeness of single-cell gene expression of gene *g* for grouping cells into two clusters *j* and *k* by the maximum likelihood principle in CALISTA. For each edge in the lineage graph, CALISTA generates a rank list of genes in decreasing values of 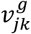. The transition genes correspond to the set of top genes in the list such that the ratio between the sum of 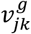 among these genes and the total sum of 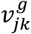 among all genes exceeds a given threshold (default threshold: 50%).

#### Pseudotemporal ordering of cells

Given a lineage progression graph among cell clusters, the third and last novel algorithm in CALISTA concerns with the pseudotemporal ordering of cells. For this purpose, we first assign a pseudotime to each cluster. If the time or stage information of the cells is provided, then the pseudotime of a cluster is set to the mode of the time/stage of the cells in the cluster divided by the largest time/stage. When the time/stage information is not available, but the starting cluster is known (e.g., from knowledge of starting cells or marker genes), we assign a pseudotime of 0 for the starting cluster. We then evaluate the sum of the cluster distances along each path in the lineage progression and identify the maximum cumulative cluster distance. The pseudotime of a cluster is given by its cumulative cluster distance to the starting cluster divided by the maximum cumulative cluster distance. Once the cluster pseudotimes have been set, we assign each cell to one of the state transition edges and compute the cell pseudotime using the maximum likelihood principle (see *Cell assignment to transition edges* below). Finally, given a developmental path in the linage progression, CALISTA provides a pseudotemporal ordering of cells that have been assigned to the edges belonging to the path.

### Cell assignment to transition edges

For pseudotemporal ordering of the cells, CALISTA first assigns cells to the edges in the lineage graph. In the following illustration, let us consider a cell *n* in cluster *k.* CALISTA allocates the cell *n* to one of the edges that are incident to cluster *k*, again following the maximum likelihood principle. For this purpose, we define the likelihood value of a cell *n* to belong to an edge pointing from any cluster *j* to cluster *k* as follows:

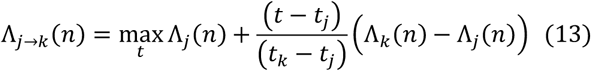

where *t_k_* denotes the cluster pseudotime label and Λ_*k*_(*n*) defines the likelihood value of the *n*-th cell to be in cluster *k*, i.e. 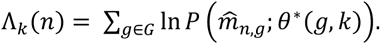 Similarly, we define the likelihood value of the same cell to belong to an edge pointing from cluster *k* to any cluster *l* by the following:

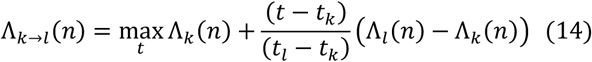

CALISTA computes all possible Λ_*j*→*k*_(*n*) and Λ*_*k*→*l*_*(*n*) lineage graph, and assigns the cell to the edge that gives the maximum of all Λ*_*j*→*k*_*(*n*) and Λ*_*k*→*l*_*(*n*) values. The pseudotime of the cell *t*(*n*) is set to *t* that gives the maximum likelihood value, as follows:

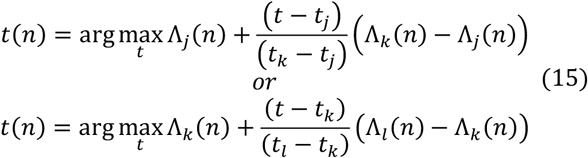

depending on the cell assignment to edges above.

### Cell ordering along a developmental path

Given a lineage progression graph, users can identify one or several developmental paths. A developmental path is defined as the sequence of connected clusters in the lineage progression graph with transition edges pointing from one cluster to the next in the sequence. CALISTA generates a pseudotemporal ordering along a given developmental path by first identifying cells belonging to the state transition edges in the path and order these cells according to their pseudotimes. Note that in defining the likelihood function for assigning cells to edges, we have assumed that the steady state probability distributions of gene expressions vary linearly between two connected clusters or states. But the result of the cell ordering does not change if we replace the linear interpolation function with any monotonic function.

### *In silico* single-cell time-stamped expression data generation

For testing the performance of CALISTA, we simulated synthetic single-cell gene expression data using the stochastic differentiation equation (SDE) model of the gene network (12 genes) governing the differentiation of central nervous system (CNS) proposed by Qiu et al. (Qiu et al., 2012, n.d.). We simulated single-cell gene expression data for 9 time points (*t_sim_* = 0, 1, 2, 4, 6, 8, 12, 16, 20) and a total of 1800 cells (i.e., 200 cells per time point) using a time step of 0.04. Prior to the sampling, we simulated the model until steady state. In order to simulate asynchronous cell differentiation, for each time point *t_sim,i_*, we took cells from random simulation times assuming a Gaussian distribution with a mean of *t_sim,i_* and a standard deviation of 0.2. The *in silico* data are included in CALISTA package (http://www.cabselab.com/calista).

