## Supplementary Figure for "CALISTA: Clustering and Lineage Inference in Single-Cell Transcriptional Analysis"

**Supplementary Notes and Figures**
**CALISTA: Clustering and Lineage Inference in Single-Cell Transcriptional Analysis**

Nan Papili Gao[1,2], Thomas Hartmann[1], Tao Fang[1], Rudiyanto Gunawan[3,*]

[1]Institute for Chemical and Bioengineering, ETH Zurich, 8093 Zurich, Switzerland

[2]Swiss Institute of Bioinformatics, 1015 Lausanne, Switzerland

[3]Department of Chemical and Biological Engineering, University at Buffalo, Buffalo, NY 14260

### Supplementary Note S1. Single-cell clustering algorithm

#### *Maximum likelihood clustering by greedy algorithm*

As illustrated in Figure 1b in the main text, the greedy algorithm for the maximum likelihood single-cell clustering in CALISTA comprises three basic steps:

*1. Initialization*. Cells are randomly assigned into $K$ clusters with equal probabilities (i.e., uniform random cell assignments).

*2. Parameter estimation.* For each cluster $k$ (between 1 and $K$) and each gene $g$ (between 1 and $G$), CALISTA obtains the parameter vector of the two-state gene transcription model $\theta^*(g,k) = \left\{\theta_{on}^*, \theta_{off}^*, \theta_t^*\right\}_g^k$ that maximizes the joint probability of obtaining the gene expression values of the cells in the cluster, as follows:

$$\theta^*(g,k) = arg\max_{\theta} \sum_{n \in N_k} \ln P\left(\widehat{m}_{ng};\theta\right) \quad (S1)$$

where $N_k$ is the set of cell indices for the cells in cluster $k$. The probability $P(\widehat{m};\theta)$ is computed using the steady state probability distribution from the two-state gene transcription model as given in Eq. (2) (assuming that random dropout events are insignificant) or using the distribution function defined in Eq. (4).

For computational speedup, as previously done in SABEC (Ezer et al., 2016), the log-probability $\ln P(\widehat{m}|\theta)$ values are pre-computed for $m$ between 0 and 200 and for a set of parameter vectors $\left\{\theta_{on}^w, \theta_{off}^w, \theta_t^w,\right\}_{w=1}^W$, producing a $W \times 201$ matrix $L$. More specifically, the ($i$, $j$)-th element of $L$ gives the log-probability $\ln P(j|\theta_i)$. The parameter set consists of all combinations from the following parameter discretization: $\theta_{on}$ =[0.5, 1, 2, 3, 4, 5], $\theta_{off}$ =[0.25, 0.5, 1,

2, 3, …, 19], and $\theta_t$ = [1, 51, 101, …, 451]. The parameter ranges in CALISTA are adapted from SABEC and selected to cover the parameter combinations for which $P(\hat{m}|\theta)$ spans a family of steady state distributions of mRNA counts, including negative binomial, Poisson, and bimodal distributions. While the number of parameter combinations ($W$ = 945) in CALISTA is much lower than that in SABEC ($W$ = 39500), our extensive testing indicated that the single-cell clustering in CALISTA performs as well as SABEC at much lower computational requirements.

By pre-computing the log-probability values, the solution to the parameter estimation problem above reduces to finding the parameter combination corresponding to the maximum element of the following vector:

$$L \times r^{k,g} \qquad (S2)$$

where $r^{k,g}$ is a $201 \times 1$ vector with its $i$-th element being the number of cells in cluster $k$ with its expression value of gene $g$ rounded to the nearest integer equaling to $\hat{m}_{n_k,g}$.

*3. Cell re-assignment.* Once the optimal parameter values for all $G$ genes and all $K$ clusters have been obtained, CALISTA reassigns each cell to its new cluster following a greedy algorithm. More specifically, CALISTA evaluates the likelihood for each cell to belong to a cluster as follows:

$$\Lambda_k(n) = \sum_{g=1}^{G} \ln P\left(\hat{m}_{n,g}; \theta^*(g,k)\right) \qquad (S3)$$

where $\Lambda_k(n)$ denotes the likelihood value of the $n$-th cell to be in cluster $k$. We reassign each cell to the cluster that gives the maximum likelihood.

Steps 2 and 3 from the algorithm outlined above are repeated iteratively until convergence, i.e. until there are no new cell reassignments.

#### *Single-cell clustering using consensus matrix*

To improve both the robustness of the single-cell clustering, CALISTA executes the iterative greedy clustering algorithm above until convergence for a user-specified number of times (by default 50), each starting with a different random initial cell assignment. The results from the independent clustering runs are combined to produce a $N \times N$

symmetric consensus matrix $C$. The (*i*,*j*)-th element of the matrix $C$ gives the number of times that the *i*-th and *j*-th cells are clustered together, as illustrated in Figure 1b. CALISTA subsequently applies *k*-medoids algorithm using the consensus matrix to obtain the final cell clustering assignment (Bhat, 2014).

#### *Choosing the number of clusters*

As an aid for choosing the appropriate number of clusters $K$ – when the number is not known *a priori* – CALISTA provides the eigengap plot based on the normalized graph Laplacian matrix $A_{N\times N}$ of the consensus matrix $C$, defined as follows (von Luxburg, 2007):

$$A = I - B^{-1/2} C B^{-1/2} \qquad \text{(S4)}$$

where $I$ is the $N \times N$ identity matrix and $B$ is the $N \times N$ degree matrix of $C$, i.e. the diagonal matrix whose elements represent the row-wise sum of the consensus matrix (i.e., $b_{i,i} = \sum_{j=1}^{N} c_{i,j}$). The eigengap plot is the plot of the eigenvalues $\lambda$ of the matrix $A$ in an ascending order. The number of clusters $K$ can be heuristically chosen based on a "gap" in the eigenvalues of $A$, where the lowest $K$ eigenvalues $\lambda_{1\ldots}$ $\lambda_K$ values are much smaller than $\lambda_{K+1}$.

#### Supplementary Note S2: *Application of CALISTA to massively parallel Drop-seq expression data*

##### *Identification of mouse spinal cord neurons activity during behavior (Sathyamurthy et al., 2018)*

The dataset comprises single-nucleus RNA-sequencing (snRNA-seq) transcriptome of 18,000 nuclei collected from adult mouse lumbar spinal cord. In the data pre-processing, we removed genes with non-zero expression values < 3 and cells in which less than 200 genes are expressed. A total of 17,784 nuclei and top 300 most variable genes were considered for further analysis (see Methods). Based on the eigengap plot, we set the number of cluster equal to 9.

##### *Analysis of peripherical blood mononuclear cells (PBMCs) (Zheng et al., 2017)*

The dataset comprises single-cell Drop-sequencing (scDrop-seq) transcriptional profiles of 68,579 PBMCs. In the data pre-processing, we removed genes with non-zero expression values < 3 and cells in which less than 200 genes are expressed. A total of 68,547 single cells and the top 300 most variable genes were considered for further analysis (see Methods). Following the clustering results of the original study, we applied CALISTA to group cells into 10 clusters.

##### *Reconstruction of developmental trajectories during zebrafish embryogenesis (Farrell et al., 2018)*

The dataset comprises single-cell Drop-sequencing (scDrop-seq) transcriptional profiles of 38,731 cells collected at 12 developmental stages (from 3.3 to 12 hours post-fertilization) during zebrafish embryogenesis. In the data pre-processing, for each time point, we removed genes with more than 90% of zero expression values and selected the top 100 most variable genes for further analysis (see Methods). In the clustering analysis of scDrop-seq from zebrafish embryogenesis (Farrell et al., 2018), we applied the following procedure:

1. Cluster cells from the final developmental stage / time $t$. The optimal number of clusters $K_t$ was chosen as the maximum $K$ among the top three eigengaps. For this dataset, we selected 23 as the number of clusters at the final time point based on the eigengap plot in Supplementary Figure S11.
2. Cluster cells from the previous stage $t-1$ with the following constraint on the number of clusters:

$$K_{t-1} \in \begin{cases} \left[\max\left(1, \left\lfloor \frac{K_t}{3} \right\rfloor\right), K_t\ \right] & if\ t > 3 \\ [1, K_t\ ] & if\ t \leq 3 \end{cases} \qquad (S5)$$

3. CALISTA automatically finds the optimal number of clusters $K_{t-1}$ within the permitted range again as the maximum $K$ among the top three eigengaps.
4. Repeat step 2 until the data from all time points are analyzed. A total of 82 clusters were identified.

After cell clustering, CALISTA selects the most informative genes as the union of the most variable genes detected at each time point and re-scales the original data accordingly (see Methods). For reconstructing the lineage progression, we considered transitions or edges only between clusters of adjacent time points, again in the order of increasing cluster distances. We added a transition edge such that each cluster has one incoming and outgoing edge, except for the initial and final clusters that have only one in out-edge and in-edges, respectively (see Supplementary File S5).

#### Supplementary Note S3: Effects of single-cell gene expression scaling on CALISTA clustering accuracy

To assess the effects of the gene expression scaling in the data preprocessing of CALISTA, we tested CALISTA clustering algorithm using *in silico* datasets as follows. First, we generated five single-cell expression datasets, each containing five cell clusters (see Supplementary File S4). In particular, we implemented Gillespie stochastic simulation algorithm to simulate the two-state model described by Peccoud and Ycart (Peccoud and Ycart, 1995). For each gene and each cluster, we randomly selected the parameter values for the two-state model from the following ranges: $0.25<\theta_n<10$, $0.25<\theta_{off}<20$ and $1<\theta_t<1500$. For these parameter ranges, the shapes of the gene distributions include bimodal, negative binomial, and Poisson distribution. We generated *in silico* single-cell mRNA counts for 50 genes and 100 single cells for each cluster, with a total of 500 cells per dataset. We followed the pre-processing and single-cell clustering procedure as described in Methods and Supplementary Note 1. We assessed the accuracy of the cell clusters generated by computing the adjusted Rand index (ARI). Comparing the CALISTA clustering results to the true clusters, the data scaling in the pre-processing step of CALISTA did not have any effects on the clustering accuracy for the five *in silico* datasets (ARI = 1 for all datasets, see Supplementary File S4).

### Supplementary Figures

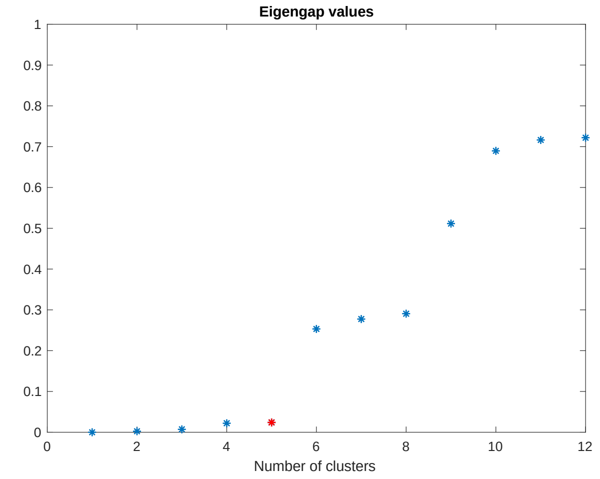


**Supplementary Figure S1. Eigengap plot of single-cell clustering analysis of Bargaje et al. dataset** (Bargaje et al., 2017)**.** Following the eigengap heuristic, the number of clusters was chosen to be five (marked in red).

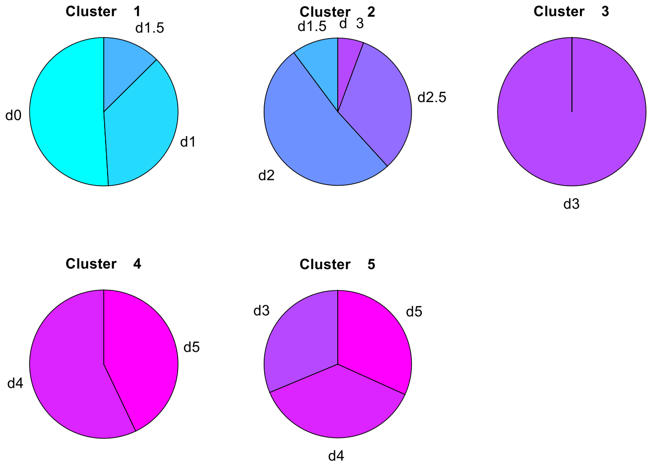


**Supplementary Figure S2. Compositions of cell clusters of single-cell clustering analysis of Bargaje et al. dataset** (Bargaje et al., 2017) The pie charts show the proportions of cells from different capture times (day 0, 1.5, 1, 2, 2.5, 3, 4 and 5), contained in each of the five clusters from CALISTA.

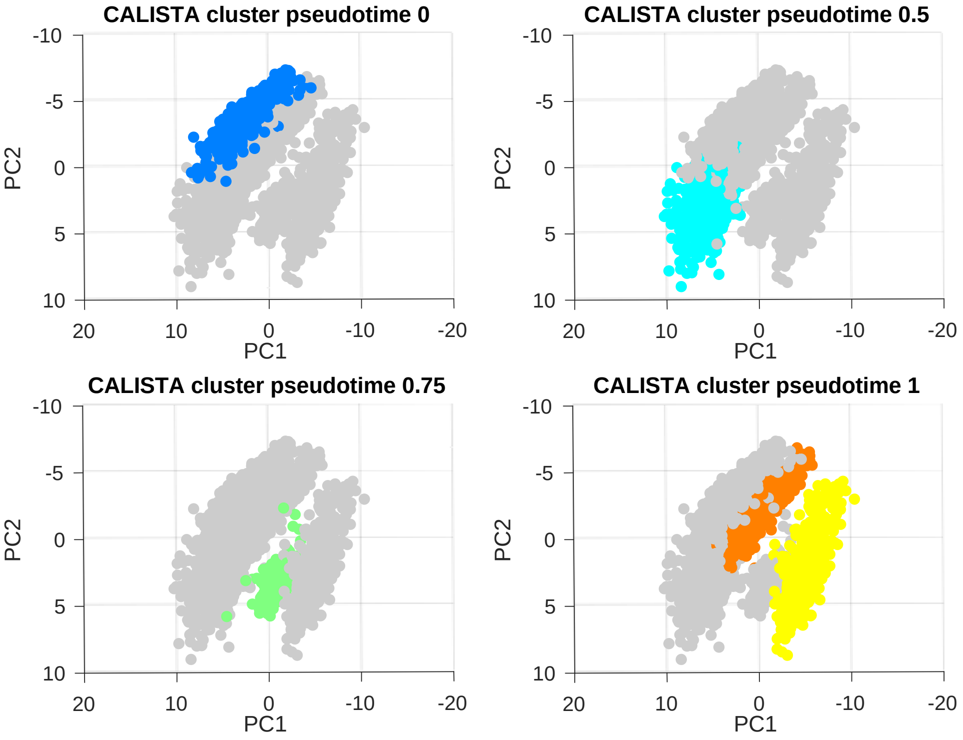


**Supplementary Figure S3. Single-cell cluster pseudotimes of Bargaje et al. dataset** (Bargaje et al., 2017) The cluster pseudotimes were set to the most frequent capture time of the cells in each cluster, normalized by the maximum capture time (see Methods).

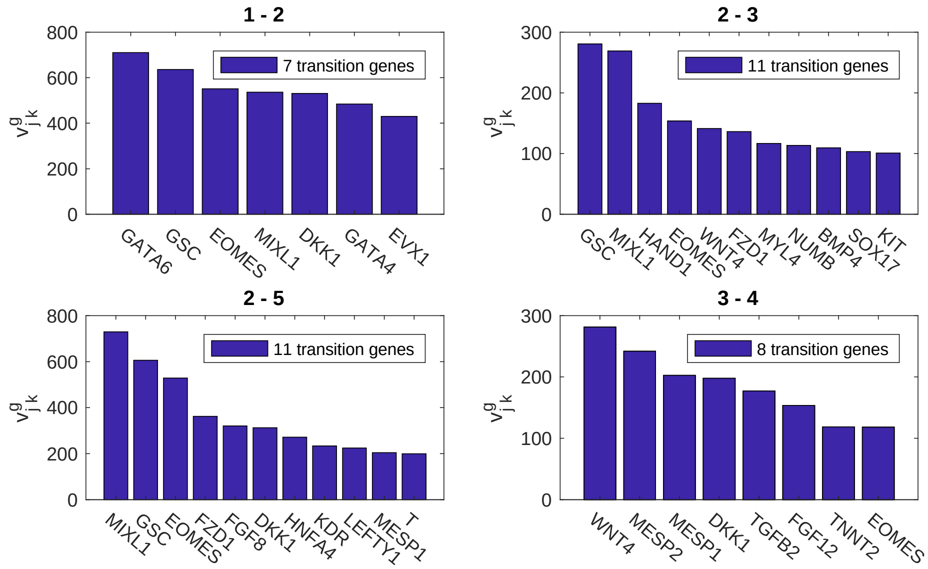


**Supplementary Figure S4. Transition genes in the lineage progression of iPSC differentiation to mesodermal and endodermal cells of Bargaje et al.** (Bargaje et al., 2017) The histograms show the transition genes sorted according to the value $v_{jk}^{g}$ (in descending order) for each of the state transition edges given in the title. The cutoff value for the inclusion of genes was chosen as the point at which the total sum of $v_{jk}^{g}$ among all genes exceeds a given threshold (default threshold: 50%).

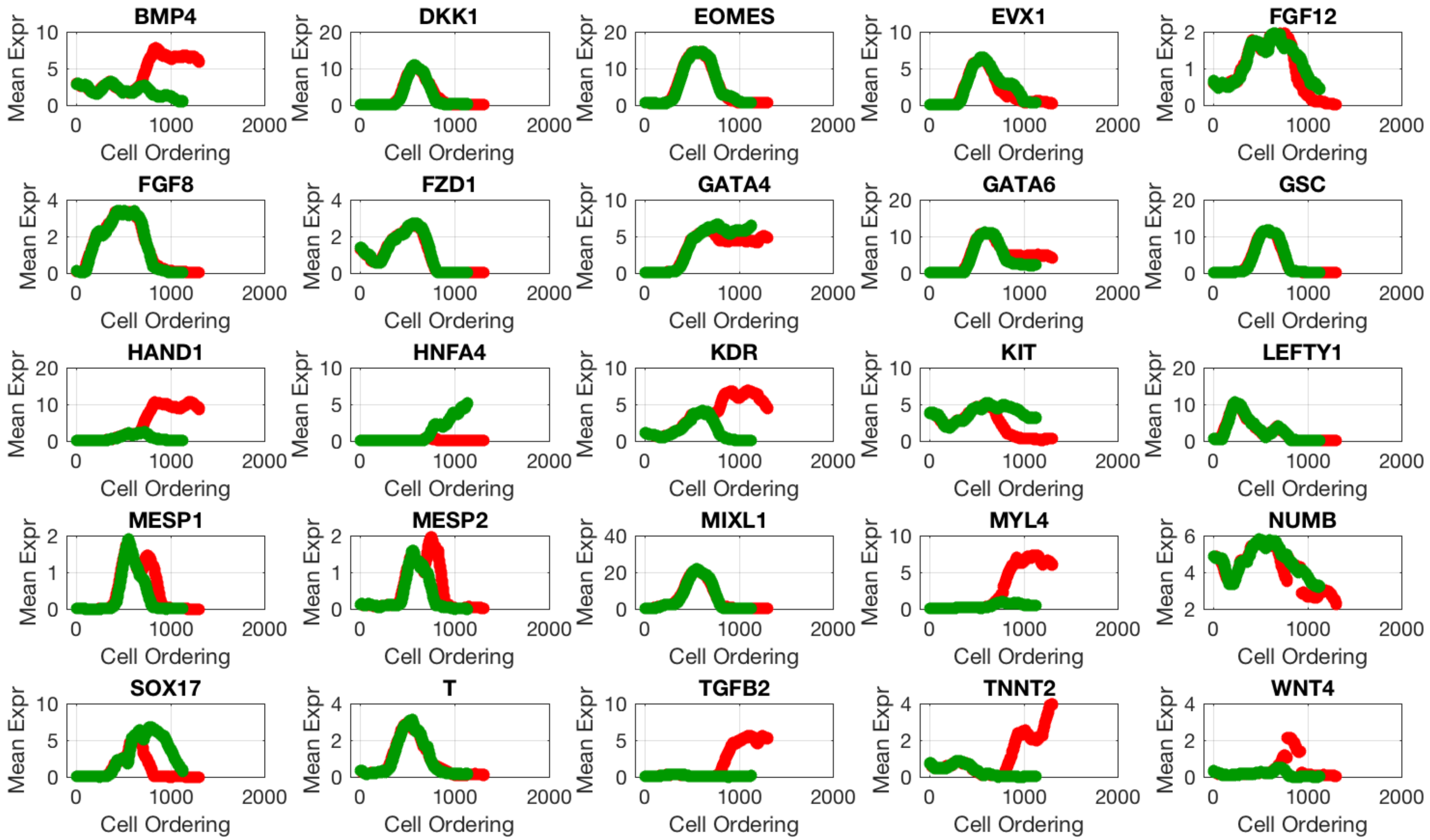


**Supplementary Figure S5. Pseudotemporal profiles of the expression of transition genes.** The subplots show the moving average expression of transition genes for pseuotemporally ordered cells along mesodermal (red) and endodermal (green) developmental paths in the single-cell transcriptional analysis of Bargaje et al. study (Bargaje et al., 2017).

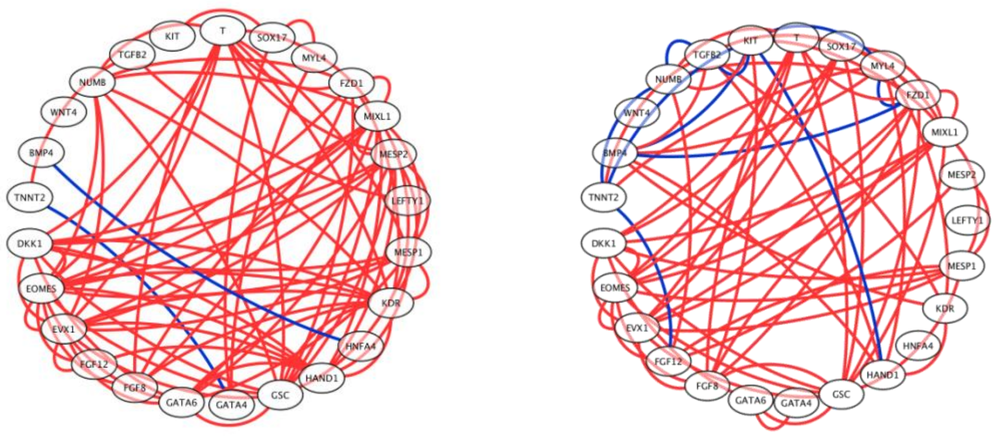


**Supplementary Figure S6.** C**o-expression networks for mesodermal and endodermal developmental paths in Bargaje et al. study** (Bargaje et al., 2017)**.** Gene co-expression network for (Top) mesodermal path and (Bottom) endodermal path. The co-expression network includes only gene-gene Pearson correlations ≥ 0.8 with *p*-value ≤ 0.01.

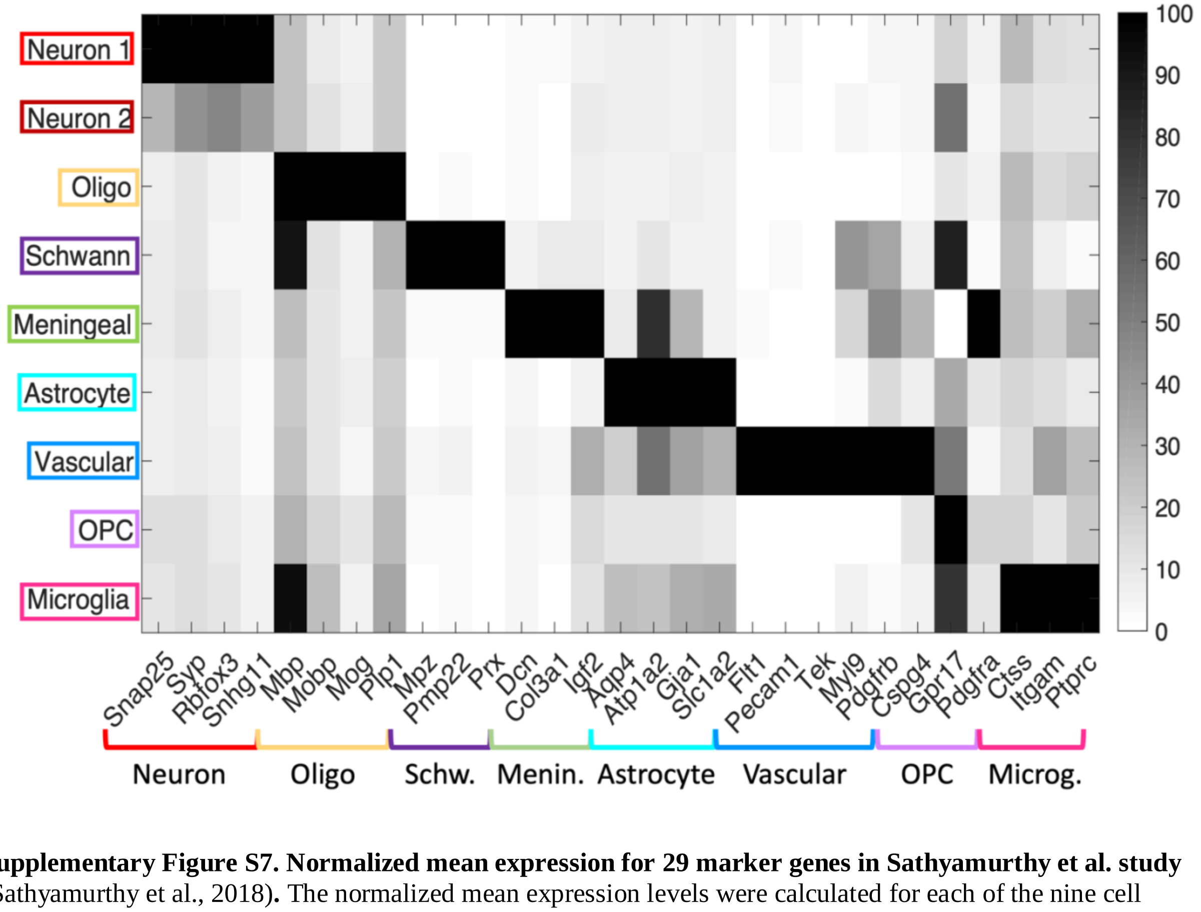


**Supplementary Figure S7. Normalized mean expression for 29 marker genes in Sathyamurthy et al. study** (Sathyamurthy et al., 2018)**.** The normalized mean expression levels were calculated for each of the nine cell clusters predicted by CALISTA.

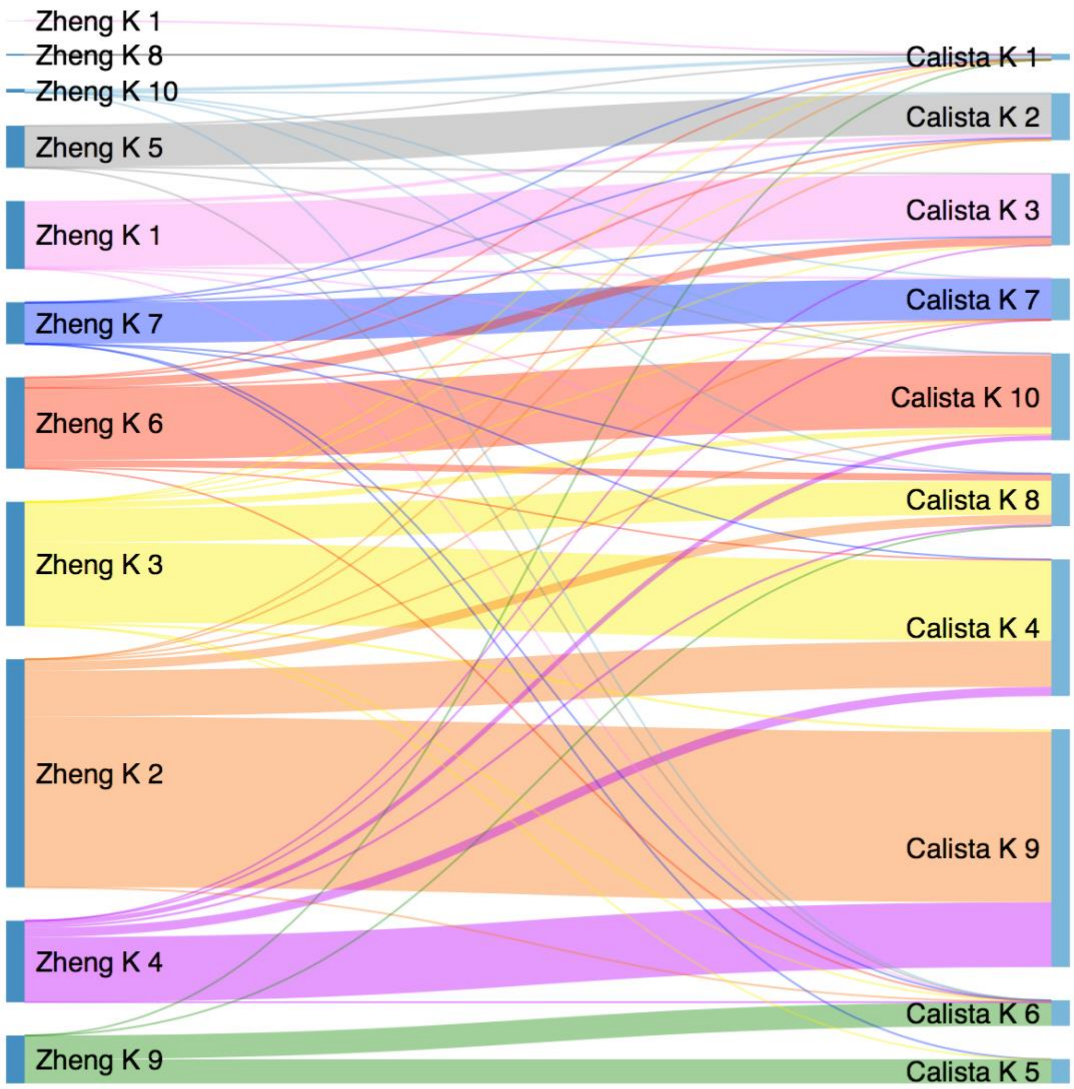


**Supplementary Figure S8. Sankey diagram comparing CALISTA clustering result with the original cell clusters in Zheng et al. study** (Zheng et al., 2017)**.** Cluster colors are taken from the original publication. Line widths are proportional to the cells in common between each pair of clusters.

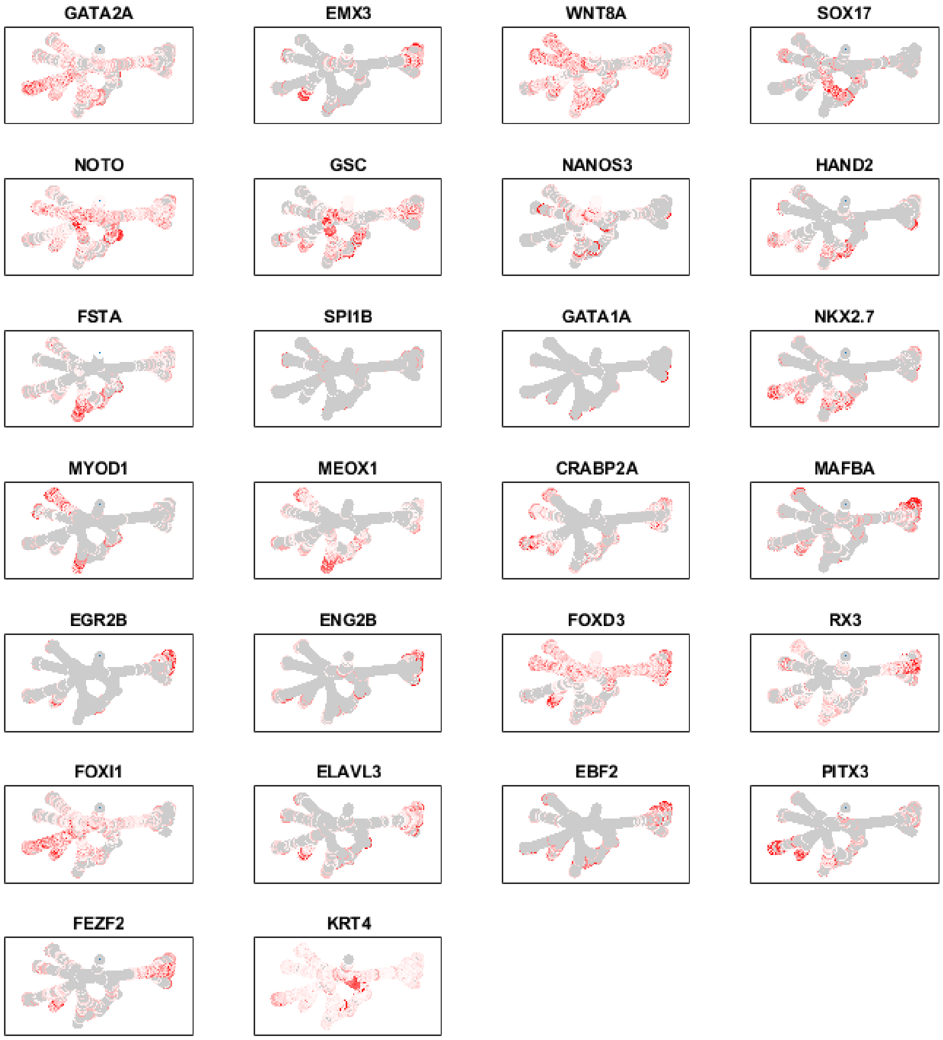


**Supplementary Figure S9. Marker gene expression during zebrafish embryogenesis.** Gene expression levels (Red = high expression, gray = no expression) were plotted along the developmental trajectories inferred by CALISTA.

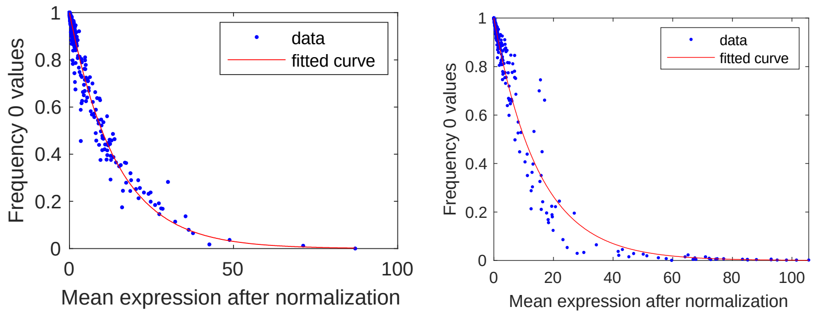


**Supplementary Figure S10. Random dropout model.** The dropout frequency is modeled as negative exponential function fitted on the data based on the gene-wise mean normalized expression versus frequency of zero-values (Kharchenko et al., 2014; Pierson and Yau, 2015). (*left*: Sathyamurthy et al. (Sathyamurthy et al., 2018), *right*: Zheng et al. (Zheng et al., 2017))

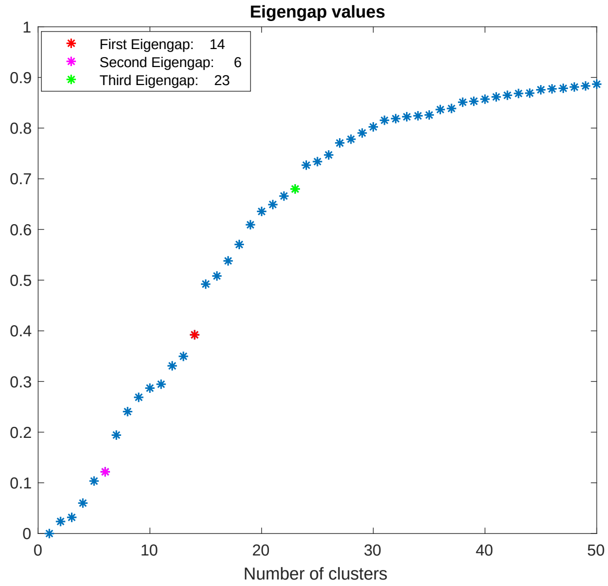


**Supplementary Figure S11. Eigengap plot of single-cell transcriptional analysis of zebrafish embryogenesis in Farrell et al. study** (Farrell et al., 2018)**.** The optimal number of clusters was chosen as the third observable gap, shown in green (K=23).
